# Ritualistic use of ayahuasca enhances a shared functional connectome identity with others

**DOI:** 10.1101/2022.10.07.511268

**Authors:** Pablo Mallaroni, Natasha L. Mason, Lilian Kloft, Johannes T. Reckweg, Kim van Oorsouw, Stefan W. Toennes, Hanna M. Tolle, Enrico Amico, Johannes G. Ramaekers

## Abstract

The knowledge that brain functional connectomes are both unique and reliable has enabled behaviourally relevant inferences at a subject level. However, it is unknown whether such “fingerprints” persist under altered states of consciousness. Ayahuasca is a potent serotonergic psychedelic which elicits a widespread dysregulation of functional connectivity. Used communally in religious ceremonies, its shared use may highlight relevant novel interactions between mental state and FC inherency. Using 7T fMRI, we assessed resting-state static and dynamic FCs for 21 Santo Daime members after collective ayahuasca intake in an acute, within-subject study. Here, connectome fingerprinting revealed a shared functional space, accompanied by a spatiotemporal reallocation of keypoint edges. Importantly, we show that interindividual differences in higher-order FCs motifs are relevant to experiential phenotypes, given that they can predict perceptual drug effects. Collectively, our findings offer an example as to how individualised connectivity markers can be used to trace a subject’s functional connectome across altered states of consciousness.

## Introduction

The uniqueness of one’s brain connectivity profile is being increasingly recognised as a ubiquitous principle of <u>connectomics</u>. Akin to the ridges and furrows that comprise our fingerprints, functional connectivity patterns derived from functional resonance magnetic imaging (fMRI) data, also known as functional connectomes (FC)^1^, have been found to be stable across a lifetime ^2^ and hold explanatory power for robust inferences at a single-subject level ^3^. Evidence has shown complex behavioural phenotypes such as cognition ^4^, demographics ^5^, traits such as fluid intelligence ^6^ or personality ^7^, and even clinical outcomes ^8^ can be reliably predicted from FCs alone. This observation has led to calls to move away from group-level inferences and towards interindividual differences prior to drawing conclusions on the generalisability of brain activity ^9,10^.

In recent years, efforts have been underway to develop the field of “brain fingerprinting” ^11^. First exemplified by Finn et al., individual subjects were shown to be readily distinguishable from a set of FCs based on their correspondence ^12^. Since then, work has demonstrated that an individual’s connectome fingerprint across sessions can be separated into signalling motifs reflecting both trait intra-subject and state-dependent inter-subject variance ^13,14^, reproducible across modalities ^15,16^, acquisition methods ^17-19^ and durations ^11,20,21^. These findings have contributed to the notion that, across mental states, there exists an “intrinsic’’ functional network architecture which is inherent to brain function and exhibits subtle variations among individuals ^22-24^. That said, it is important to consider that these fingerprints of brain organisation might not just be limited to the spatial organisation and independence of FC traits, but likely also to their temporal quality ^25^. Spatiotemporal dynamics have been suggested to provide a ‘common currency’ for mental and neuronal states ^26^, with neural processing being organised across timescales and increasing along the cortical hierarchy of information processing ^27^. According to this view, shorter timescales in sensory areas facilitate the rapid detection and encoding of dynamic stimuli, which are subsequently integrated by the slower dynamics of associative areas over longer time frames ^28,29^.

Much work has been concerned with understanding how such inherent connectivity might be differentially altered according to a particular individual or mental state ^15,24,30^. However, there is little evidence bridging these lines of research, particularly quantifying the variance associated with a subject versus the brain state under which it is examined ^31^. Altered states of consciousness (ASCs) induced by serotonergic psychedelics may provide a new means by which to probe the interdependency of unique spontaneous brain activity and the functional brain organisation during a transient disruption. Compelling evidence reveals agonism of the 5-HT_2A_ receptor by hallucinogens holds a central role in shaping drug-induced phenomena ^32^ and potentially the immanence of FC. Whole-brain modelling has implicated 5-HT_2A_ receptor distribution in shaping brain dynamics ^33^ whereas its stimulation enhances the temporal diversity of brain activity ^34^.

From this stem shifts in functional coupling between large-scale networks, ultimately diminishing integrative processing across major brain networks ^35^. Within a hierarchical predictive processing framework, these outcomes are hypothesised to be linked to the subjective experience via the decreased confidence in priors encoded by functional hierarchies ^36^. Accordingly, it may therefore be the case that stimulation of 5-HT_2A_ receptors could perturb behaviourally relevant brain fingerprints otherwise residing within high-order functional networks ^37^. Indeed, classical psychedelics have been speculated to be potential therapeutic interventions by improving symptomatology through rebalancing aberrant brain states ^36,38,39^.

A relevant practice that is purported to achieve a communal ASC is the religious use of the psychedelic brew ayahuasca. Devised from a combination of two different plant sources, the vine *Banisteriopsis caapi* and *Psychotria viridis*, ayahuasca produces a profound change to subjective experience, comprising a diffuse state of cognition alongside complex changes to self-referential awareness, perception, cognition, and mood ^40^. Whereas *Psychotria viridis* is a rich source of the potent 5-HT_2A_ agonist N,N-dimethyltryptamine (DMT), *Banisteriopsis caapi* contains monoamine oxidase inhibitor (MAOi) β-carbolines such as harmine, harmaline, and tetrahydroharmine, serving to promote the bioavailability of DMT ^41^ Historically, ayahuasca is used by syncretic religions such as *Santo Daime* to achieve personal insight, intimacy and spiritual development ^42^ Members of the congregation drink ayahuasca (termed *Daime*) communally in a ceremony referred to as the “works” (*trabalhos*). These are collective endeavours performed by members of the congregation consisting of alternating periods of song, dance, and attentive silence. Providing a formalised type of set and setting, members follow a prescribed mental state with which to engage their symbolic and religious framework (*doctrina*) ^43^. This religious use of ayahuasca might therefore provide a useful means by which to investigate the orthogonality between trait and state FC under conditions in which an individual transitions from a normal, waking state of consciousness to a shared altered state.

Here, we sought to understand how the inherency of a subject’s FC might alter under the altered state of consciousness induced by the religious consumption of ayahuasca brew. Replicating the brain fingerprint framework in Santo Daime members, we characterised changes to both static and dynamical connectome identifiability at peak drug effects. Furthermore, we explored how changes to an individual’s underlying functional connectivity might subsequently help explain aspects of their subjective experience.

## Results

Experienced members of Santo Daime were enrolled in a fixed-order, within-subject, observational study. Baseline resting-state fMRI was followed 1 day later with a second fMRI scan 90 minutes after communal intake (i.e., peak effects). The study also entailed pharmacokinetic sampling, questionnaires pertaining to retrospective drug effects and aspects of “*work*” during resting-state (see Methods). Of the 24 patients recruited, 3 were excluded due to excessive fMRI head motion. Demographic information pertaining to the imaging sample can be found in Table S1.

### Acute effects of Ayahuasca

Ayahuasca intake was associated with increased ratings on all (sub)dimensions of the 5D-ASC (*M =* 45.29 – 3.09, *t(20)* = 7.29 – 3.90, *p* < 0.001, *d* = 1.59–0.85), and for the EDI (*M* = 35.80, *p* < 0.0001, *t(20)* = 7.15, *d* = 1.56). Tandem pharmacokinetic analyses also demonstrated serum concentrations of DMT (the principal psychoactive constituent of ayahuasca) were significantly greater than zero at both 60 (*M* = 18.36 ng/ml, *t*(17) = 4.82, *p* < .0001, *d* = 1.14) and 160 minutes (*M* =7.30 ng/ml,, *t*(18) = 5.16 *p* < .0001, *d* = 1.18) after intake.

During resting-state acquisition, participants reported significantly more internal singing under ayahuasca (*W*= 58, *Z*= 2.25, *p*= 0.0261, *d* = 0.63). Session recollection nor engagement in meditation significantly differed between conditions (*p* > 0.05). A full characterisation of all inventories and serum alkaloids can be found in the supplementary materials.

### Quantifying whole-brain fingerprints

Connectome fingerprinting provides a window into the “uniqueness” of one’s functional connectivity ^11,25,44^. This approach stems from the simple assumption that a FC should hold greater similarity between test-retest scans of the same subject than between different subjects ^12^. By computing an “Identifiability matrix” we can extrapolate for a subject both *inherent* elements of functional connectomes (I_self_) and other more *shared* qualities (I_others_) which are otherwise state-dependent (see Methods, Fig.1). These measures can be summarised as a differential identifiability score (I_diff_) or, the difference between each subject’s FC self-similarity against the other subjects’ FCs. As a first pass, we explored changes to whole-brain fingerprints and their dynamical counterparts. We derived measures of identifiability for static FCs (Fig.2) and their dynamic equivalents (Fig.3) by replicating our analyses across increasing window lengths (70, 140, 210, 280 288, and 576s, with a fixed sliding window step of 7.2s). In each case we also provide success rate (SR)^12,13^ as a supplementary assessment of identifiability.

**Figure 1.**
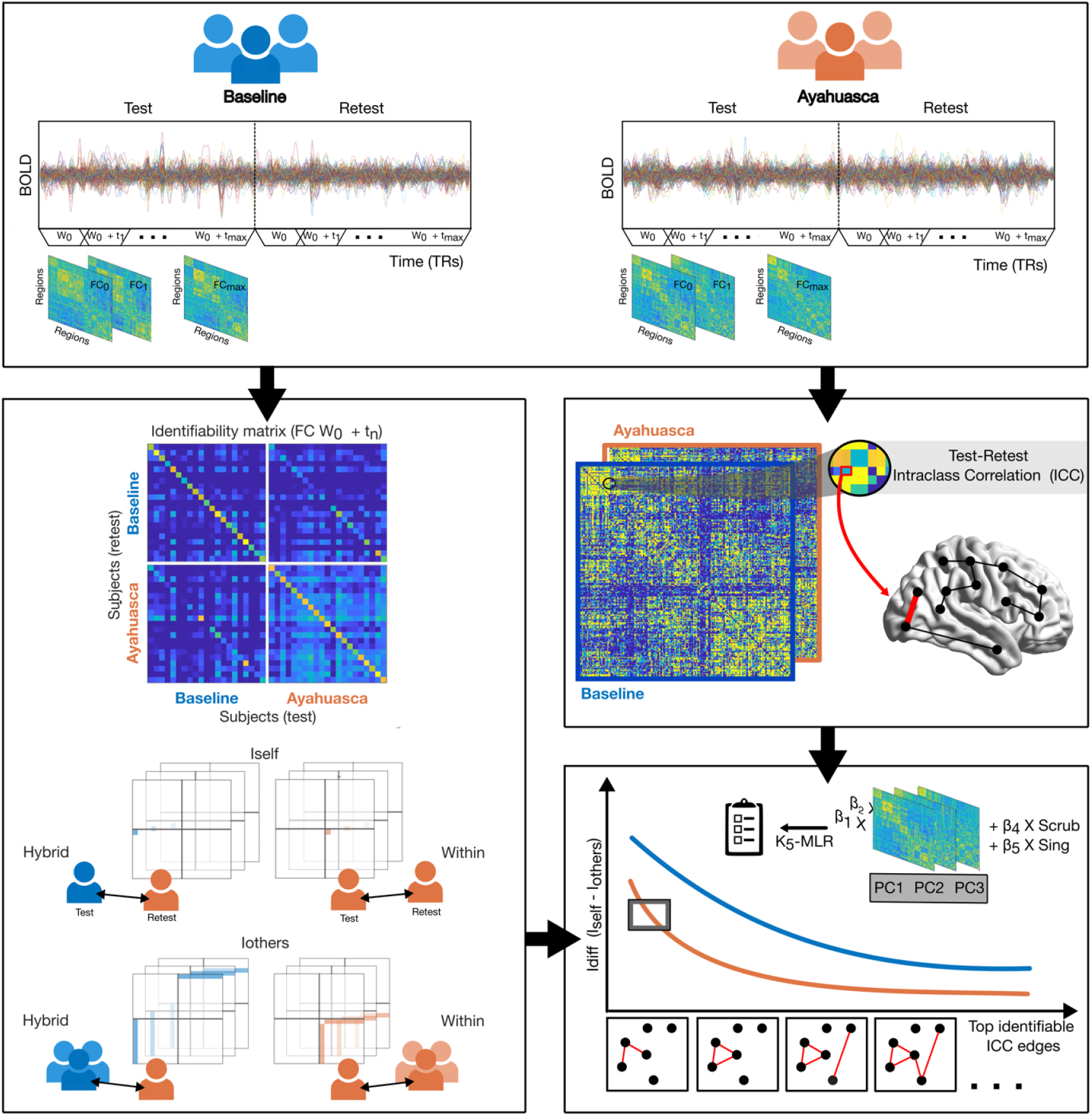
Connectome fingerprinting workflow. First, each subject fMRI timeseries is split into test vs retest halves. For all subjects and sessions, dFC frames are computed for increasing temporal windows until t_max_ is reached. Connectome fingerprints can next be calculated as the similarity in functional connectivity for all combinations of FC test vs retest (within condition and between), yielding an identifiability matrix^19^ per timescale (left). Each colour matched block reflects identifiability *within* a condition whereas colour mismatched (“hybrid”) blocks represent the distance of each subject’s identifiability *between* conditions. This object allows us to compute for each subject: I_self_ (represented by each diagonal element) denoting their similarity to oneself and I_others_ (represented by each off diagonal) representing their similarity to others, for both *within* and *between* conditions. In parallel, we can assess the fingerprinting value of specific edges per condition and timescale by calculating their intraclass correlation coefficient (ICC, right). We can next rank edges according to their ICC and iteratively calculate a compound measure of I_self_ and I_others_ (I_diff_). This allows us to examine how edges “driving” one’s fingerprint evolve under Ayahuasca. Lastly, we can assess their experiential relevance by fitting an iterative multi-linear model comprising PCA-derived principal components (PCs) of their functional connectivity as predictors of interest. Decomposing their signal into components ranked according to explained variance (PC1-3), the relevance of cohort-level (high-variance – PC1) and individual-level functional connectivity information (low-variance – PC3) is simultaneously assessed, while accounting for motion and “working”. At each step, model performance and generalisability are measured using k-fold (K_5_) cross validation, yielding an optimal edge cut-off.

**Figure 2.**
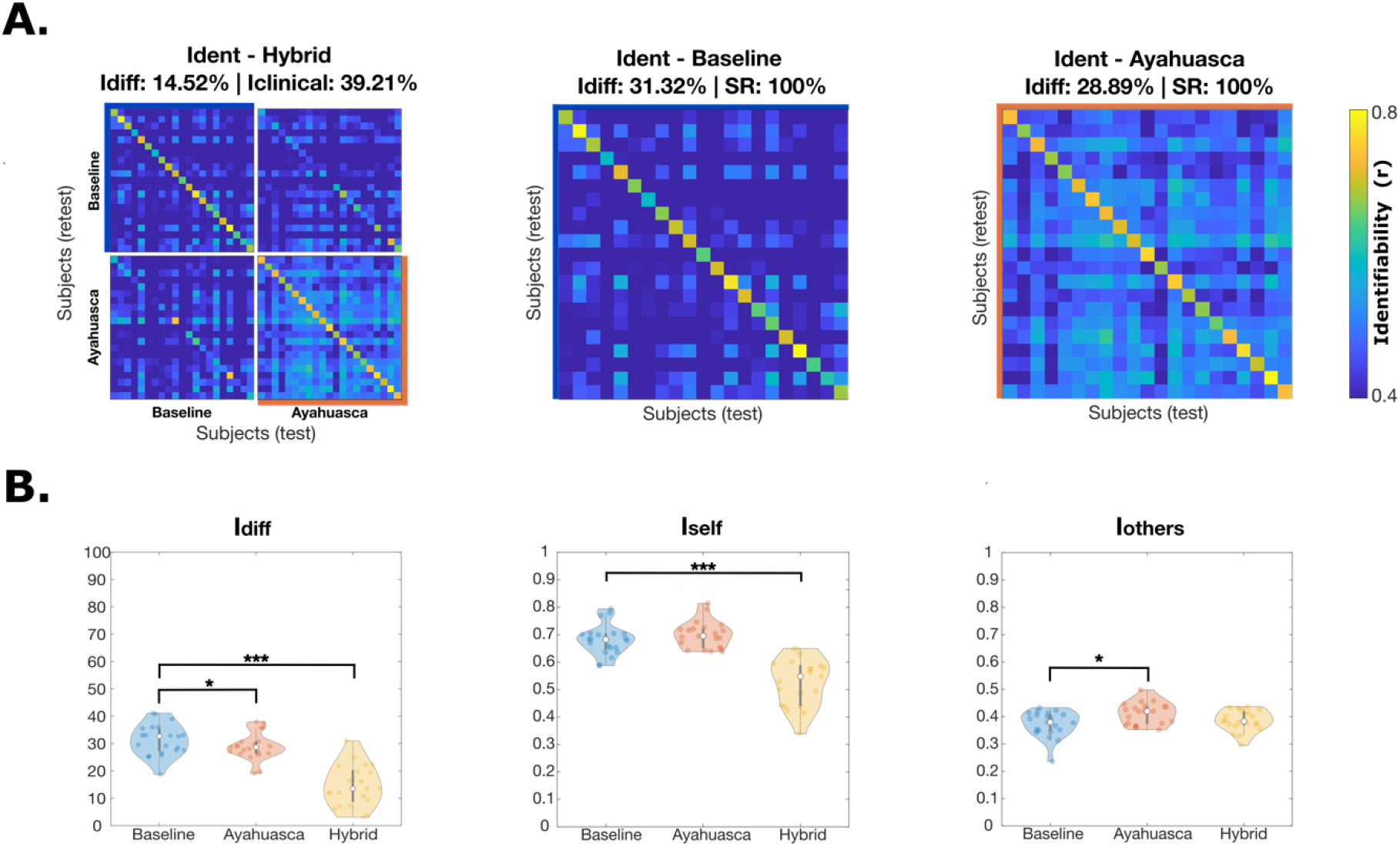
Whole-brain measures of static identifiability. **(A)** shows the identifiability matrix (far left) at T_max_ with corresponding “standard” identification matrices for each condition expanded on the right. From hybrid off-block elements one can also define the I_clinical_ for a participant as the average similarity of the individual connectome of a subject with respect to the baseline. For all, differential identifiability (I_diff_) values and success rates (SRs, where applicable) on top also provide complementary scores of the fingerprint level (see Methods). (**B)** Violin plots highlighting the difference of each identifiability metric (I_diff_, I_self_, I_others_) between conditions. Hybrid counterparts are also presented in respect to baseline. The boxplot extends from the lower to upper quartile values with a line at the median; the whiskers extend from the upper/lower quartiles up to 1.5 times the interquartile range. Subjects are represented with single points. Two-tail significance is denoted as follows: *p* < 0.05*, p < 0.01**, *p* < 0.001***.

**Figure 3.**
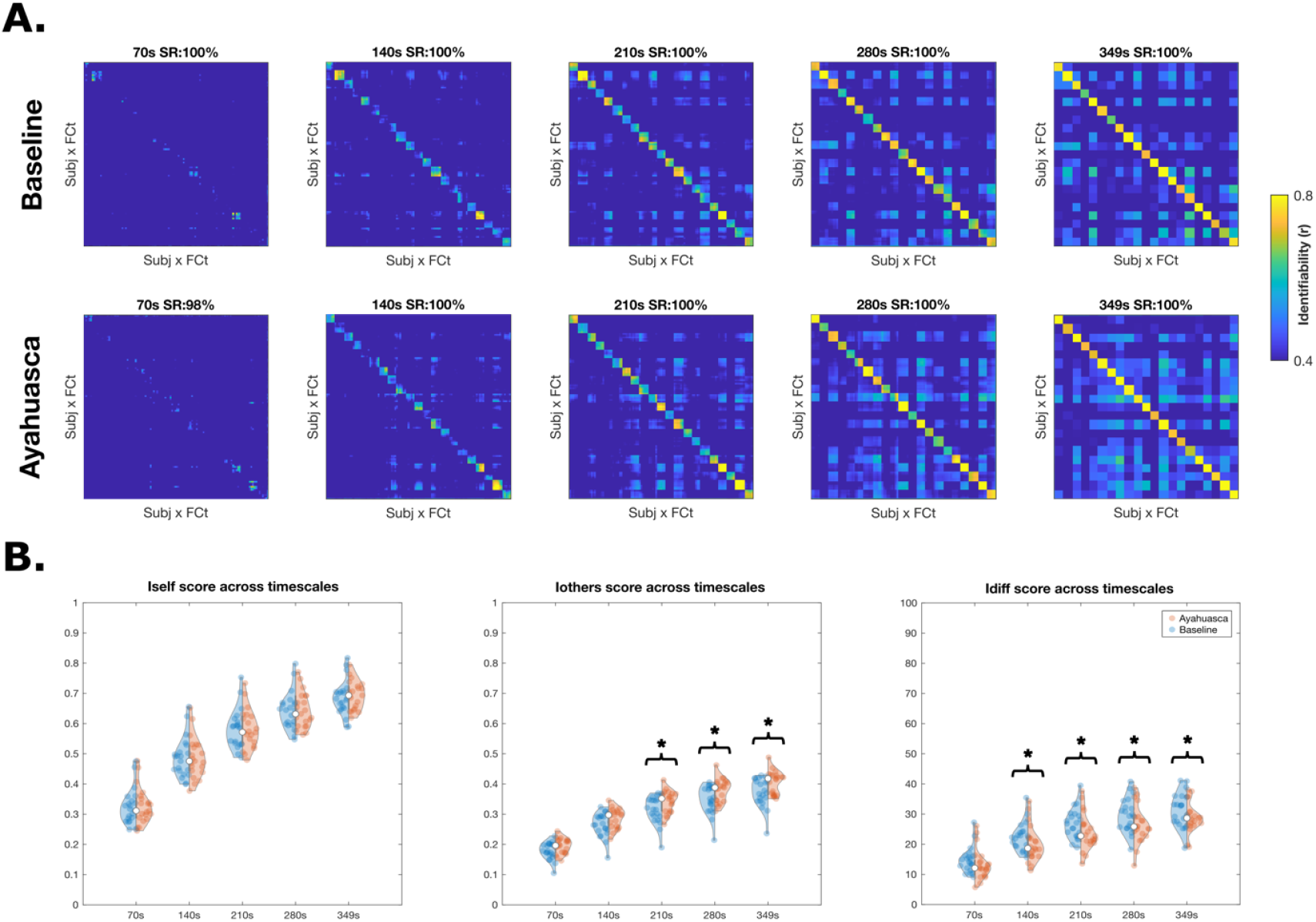
Whole-brain measures of dynamic identifiability. **(A)** Dynamic identifiability matrices at five different window lengths (70, 140,210,280 and 349s) for each condition. The dynamic differential identifiability (I_diff_) values and success rates (SRs) on top of each matrix provide two complementary scores of the fingerprint level of the dataset across temporal scales (see Methods). (**B)** Violin plots highlighting differences in each identifiability metric (Idiff, Iself, Iothers) per timescale. Each boxplot extends from the lower to upper quartile values with a line at the median; the whiskers extend from the upper/lower quartiles up to 1.5 times the interquartile range. Subjects are represented with single points. Two-tail significance is denoted as follows: *p* < 0.05*, p < 0.01**, *p* < 0.001***.

#### Static identifiability

As depicted in Fig.2B, sign-rank testing revealed the differential identifiability (I_diff_) of each participant was significantly diminished under ayahuasca (*W* = 53, *Z* = −2.17, *p* = 0.0298, *d* = 0.35). If we examine its constituents, this effect was driven by a significantly increased I_others_ score (*t*(20) = 2.72, *p* = 0.0131, *d* = 0.59). In other words, participant connectomes significantly mirrored one another’s under ayahuasca, depicted by the saturation of off-diagonal elements under Ayahuasca (Fig.2A). Remarkably, subjects continued to retain high I_self_ (*p* > 0.05) and SR scores, reflecting a preserved idiosyncrasy under Ayahuasca. It should be noted that each condition’s I_diff_ and SR scores were also significantly greater than their null equivalents following permutation testing (*p* = 0.001, see Fig.S1).

We then examined how dissimilar might the constituents of a subject’s fingerprint be under ayahuasca. Hybrid equivalents of our identifiability matrices (see Fig.1, Methods) enabled us to derive the “distance” of each subject’s score from baseline. Doing so, we identified a greater dissimilarity between a subject’s functional connectome under Ayahuasca and baseline, with both I_selfHybrid_ (*t*(20) = −8.67, *p* < 0.0001, *d* = 1.89 and I_diffHybrid_ (*t*(20) = −7.94, *p* <0.0001, *d* = 1.74) significantly reduced. In other words, while a subject’s similarity remains the same, their fingerprint makeup is reconstituted. This can be visualised as the faded diagonals in the off block “Hybrid” elements of Fig. 2A.

#### Dynamic identifiability

Might specific timescales of neural processing account for these global differences? Repeating this previous analysis across increasing window size, reveals an equivalent pattern. As per prior work ^25^, dynamic (I_diff_) increased steadily with longer window lengths (Fig.3A) as a by-product of the increasing number of timepoints for dFC computation with early dynamical fingerprints (designated by clear diagonal elements) arising at shorter temporal intervals.

Replicating our analyses across each temporal scale, we observed the equivalent patterns of change. As shown in Fig.3B, I_diff_ was significantly reduced under Ayahuasca in a temporally selective manner (across 140s-349s; max.220s: *W* = 51, *z* =-2.31, p = 0.0250, *d* = 0.41). This effect was partly accounted for by Iothers increasing across select frames (210-349s max.349s: *t* (20) = 2.72, *p =* 0.0131, *d* = 0.59). Once more, I_self_ was found to remain stable at all timescales (*p* > 0.05). For all windows, I_selfHybrid_ was significantly reduced under ayahuasca (max.210s: *t*(20) = 9.39 *p <* 0.0001, *d = 1*.*74)*, as well as I_diffHybrid_ (max.140s: *t*(20) = 9.81 *p <* 0.0001,*d* = 2.16).

### Select edges mediate reductions in connectome identifiability

Identifying global changes to each subject’s connectome fingerprint under ayahuasca, we then sought to understand their spatiotemporal profiles. To do so, we applied an edgewise ICC to investigate the fingerprinting value of edges (see Methods, Fig.1) pertaining to canonical resting-state networks (RSNs).

#### Static connectomes

We observed global reductions in ICC scores (*W* = 92864811, *z* = −7.5797, *p* <0.0001, *d* = 0.05) under Ayahuasca. While this suggests a connectome-wide drop of temporal stability, individual RSNs have varying levels of importance for fingerprints and may be differentially affected. For example, note the poor identifiability of the limbic (L) network (Fig.4A). Focusing on network properties (Fig.4B), within-network analyses revealed significant reductions in stability for the ventral attentional (VA. *W* = 8414, *z* = −4.05, *p* = 0.0014, *d* = 0.27) and converse increases for the dorsal attentional network (DA. *W* = 30021, *z* = 3.29, *p* = 0.0283, *d* = 0.20) under ayahuasca. In contrast, inspection of between-network pairs reveals reductions in stability were primarily attributable to the visual (VIS) and VA functional subsystems (*W* = 36803, *Z* = −11.54, *p* <0.0001, *d* = 0.49). Within these, certain edge pairs such as SM-L (*W* = 25535, *Z* = 5.37, *p* <0.0001, *d* = 0.26) and VIS-DA connectivity (*W* =122508, *Z* = 4.18, *p* = 0.0008, *d* = 0.14) exhibited greater stability under ayahuasca.

**Figure 4.**
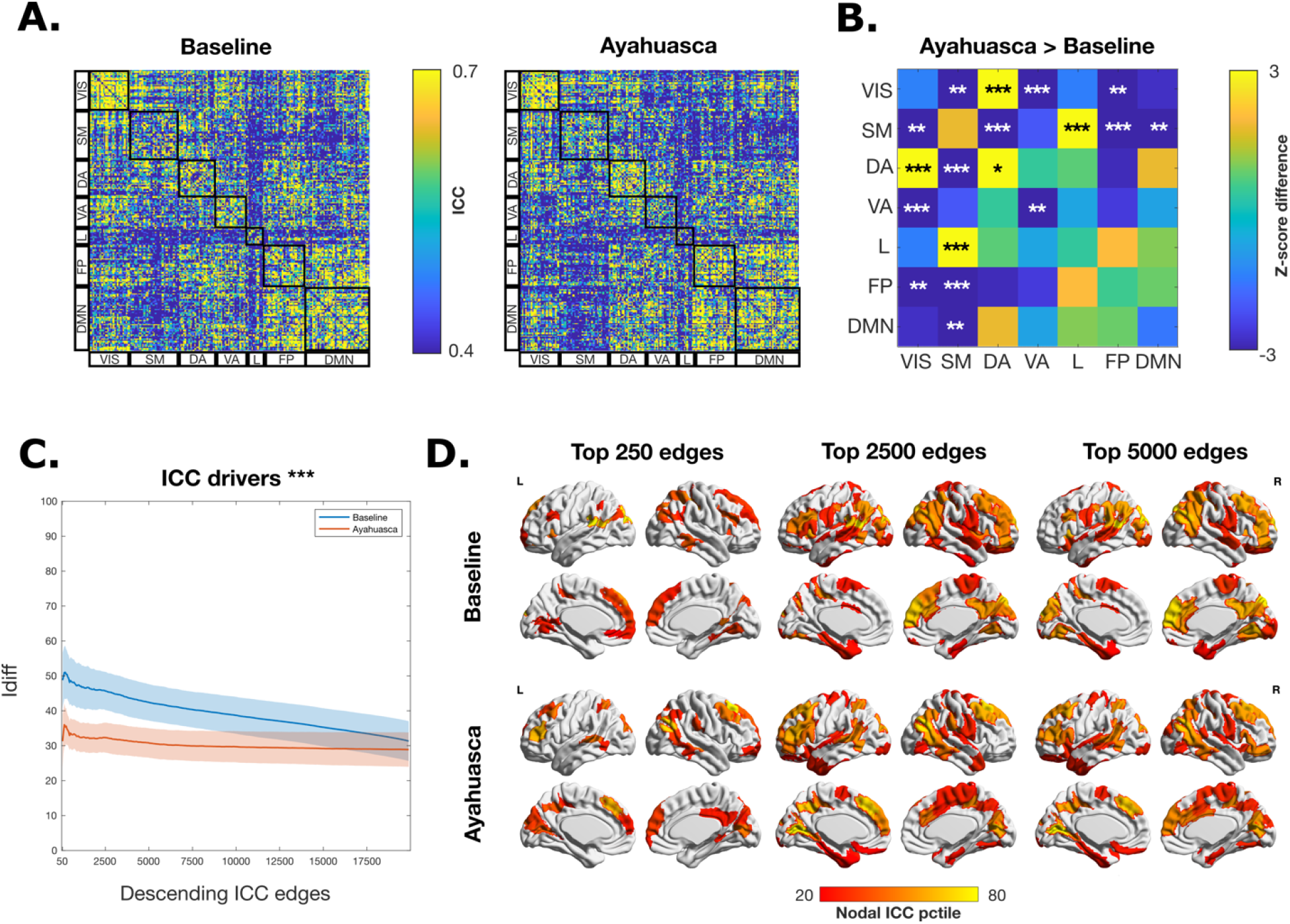
Spatial specificity of connectome fingerprints. **(A)** Edgewise intraclass correlation (ICC) matrices per condition. The ICC matrices are shown thresholded at 0.4. All 7 functional networks as defined by Yeo et al (see Methods) are highlighted by black boxes: VIS = visual network; SM = somatomotor network; DA = dorsal attentional network; VA = ventral attentional network; L = limbic network; FPN = fronto-parietal network; DMN = default-mode network. **(B)** Differences in network ICC values between conditions. For each condition, ICC edgewise scores are averaged across Yeo functional networks and compared using two-tail sign-rank testing. Approximated z-scores are then extrapolated and plotted for ease of visualisation. **(C)** Identification of top fingerprinting edges. I_diff_ scores were obtained by iteratively calculating identifiability matrices for each condition, ranked according to those contributing the most to baseline identifiability (as per ICC values). Lines represent means, with shading reflecting the standard deviation of I_diff_ across subjects at each step. **(D)**. Nodal strength (sum across unthresholded ICC matrix rows) across subsets of top fingerprinting edges per condition. For each render percentiles are shown (from 20th to 80th percentile). For all plots, two-tail significance is denoted as follows: *p* < 0.05*, p < 0.01**, *p* < 0.001***.

Given that subsets of highly synchronous edges are important contributors to normative connectome fingerprints ^11,13,25,44^, we investigated how they might shift in importance under Ayahuasca. Ranking baseline edges from most to least stable, we recalculated each subject’s identifiability 50 edges at a time. Figure 4C shows that, while baseline, or “normative” identifiability can be maximised within 250 edges, the contribution of these edges to fingerprinting drops markedly under ayahuasca (I*diff. t*(249) = −10.12, *p* <0.0001, *d* = 2.38). Therefore, edges otherwise normally “driving” a subject’s identifiability are no longer significant contributors when under the influence. Rather, a reconstitution of edge importance becomes apparent when examining their nodal equivalents (Fig.4D). One can notice connections implicated in hubs pertinent to DA, VA and SM networks are instead primarily replaced by those pertinent to the DMN.

#### Dynamic connectomes

We then examined differences in spatial ICC patterns as a function of time by repeating our analysis across each timescale. As window length increases, one can note different networks appearing at different rates, such as the VIS network at shorter intervals or the DMN at slower scales (Fig.5A). This gradient highlights the varying temporal prerequisites of RSN fingerprints ^25^Our ICC analyses revealed global reductions in dFC stability across all measured timescales (max.70s : *W* = 79109036 *z* = −24.55, *p* <0.0001 *d* = 0.18) under ayahuasca.

**Figure 5.**
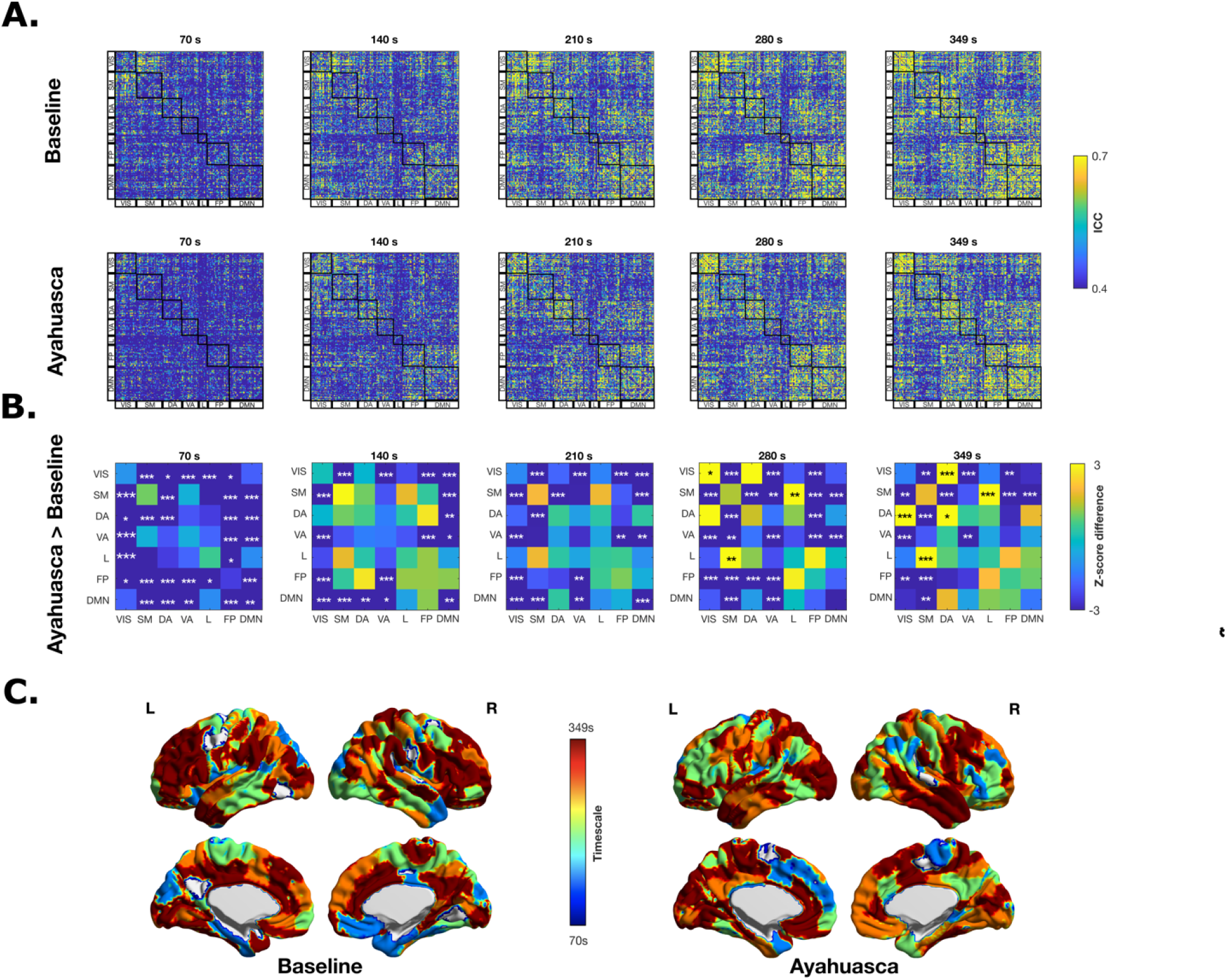
Temporal specificity of connectome fingerprints. **(A)** Edgewise intraclass correlation (ICC) matrices per condition at each timescale. The ICC matrices are shown thresholded at 0.4, the cut-off for a reliable ICC score ^72^. All 7 functional networks as defined by Yeo et al (see Methods) are highlighted by the black boxes: VIS = visual network; SM = somatomotor network; DA = dorsal-attention network; VA = ventral-attention network; L = limbic network; FPN = fronto-parietal network; DMN = default-mode network. **(B)** Differences in network ICC values across timescales. For each condition and per window, ICC edgewise scores are averaged across Yeo functional networks and compared using two-tail sign-rank testing. Approximated z-scores are then extrapolated and plotted for ease of visualisation. Lighter hues reflect increases in ICC values under ayahuasca whereas darker ones reflect diminishments under ayahuasca. **(C)** Temporal peaks of nodal stability. Maximum values across temporal profiles at each brain node are overlaid onto a brain render to map the time scales of human brain fingerprints. The maximum value for each brain node was derived from ICC nodal strength values (sum across ICC matrix rows) at each window per condition. For all plots, two-tail significance is denoted as follows: *p* < 0.05*, p < 0.01**, *p* < 0.001***.

As shown in Figure 5B, network-based analyses revealed diffuse changes to the stability of dynamic functional connectivity under ayahuasca (for a full characterisation see tables S5.1-5). Novel reductions in within-network edge stability were identified across increasing windows of time for: the DMN (max.210s: *W* = 171684, *z* = −5.89, *p* <0.0001, *d* = 0.18); VA (max.280s: *W* = 6549, *z* = −5.31, *p* <0.0001 *d = 0*.37) and DA networks (max.70s: *W* = 8250, *z* = −4.39, *p* = 0.0003, *d* = 0.245). Contrarily, VIS network edges exhibited greater stability at 280s (*W* = 48931, *z* = 3.52, *p* = 0.0122, *d* = 0.13). In parallel, reductions in between-network edge stability populated all scales. This attenuation could be primarily ascribed to edges involved in between-network SM and VIS connectivity (max.VIS-SM (70s): *W* = 82429, *z* = −13.634, *p* <0.0001, *d* = 0.47). Furthermore, previously identified static increases in SM-L connectivity stability were found to be time-dependent (280s. *W* = 18936, *z* = 3.62 p <0.0001 *d* = 0.16). We also examined whether changes to dynamic connectivity of different RSNs (see supplementary materials) might also help explain changes to the topography of edge stability. In this regard, while between-network reductions in functional connectivity variability was observed, no clear association could be ascertained (see Fig.S4).

We next asked whether altered fingerprint dynamics under ayahuasca could also be reflected at a regional level of brain organisation. Identifying each region’s ICC maximum, we summarised their temporal optimums as a brain render (Fig.5C). Typically, transmodal regions comprising association cortices, “peak” at longer temporal windows whereas unimodal regions such as primary sensory areas, arise early on ^25^. While this was the case at baseline, this temporal gradient shows an inversion effect following intake, best demonstrated by regions such as the prefrontal cortex peaking early on or vice versa for unimodal areas such as the visual cortex.

### Connectome fingerprints are predictive of perceptual drug effects

We lastly performed an exploratory analysis investigating the behavioural relevance of connectome fingerprints. We hypothesised that highly identifying edges under ayahuasca could also predict meaningful aspects of a subject’s subjective experience. To retain subject-level differences in edge connectivity, we built an iterative multilinear model approach comprising PCA components of subsets of edges as predictors of interest for our subjective effect measures. A K-fold cross validation revealed that peak predictive performance for the 5D-ASC dimensions Visual Restructuralisation (VR) and Auditory Alterations (AA) was achieved using the top 3000 most stable edges (see Figure S5.), with predictive performance for all other outcome measures being no better than the null model.

Other than three edge-based PCA components, each model also comprised two other predictors: scrubbing and internal singing. We found that of all PCA components, only PCA3 significantly increased the predictive power of the model fo**r** VR (F(4,20) = 1.98; R2 = 0.45; p = 0.0396; β = −5.59) and AA (F(4,20) = 4.8; R2 = 0.70; p = 0.0016; β = −2.17. Together, this finding might reflect the fact that subsets of edges are predictive of each dimension only when considering individual differences in functional connectivity. Edges most implicated (top 80^th^ percentile) were primarily found in FPN, DMN and DA hubs as well as the regions pertaining to the VIS (Fig. 6C). This distribution is consistent with previous literature indicating that sensorimotor networks exhibit lower inter-subject functional connectivity variability than associative networks ^45^. As a precaution we also examined whether the motion (nScrub) or shared behaviour (Singing) were relevant predictors, finding no significant contribution to model performance.

**Figure 6.**
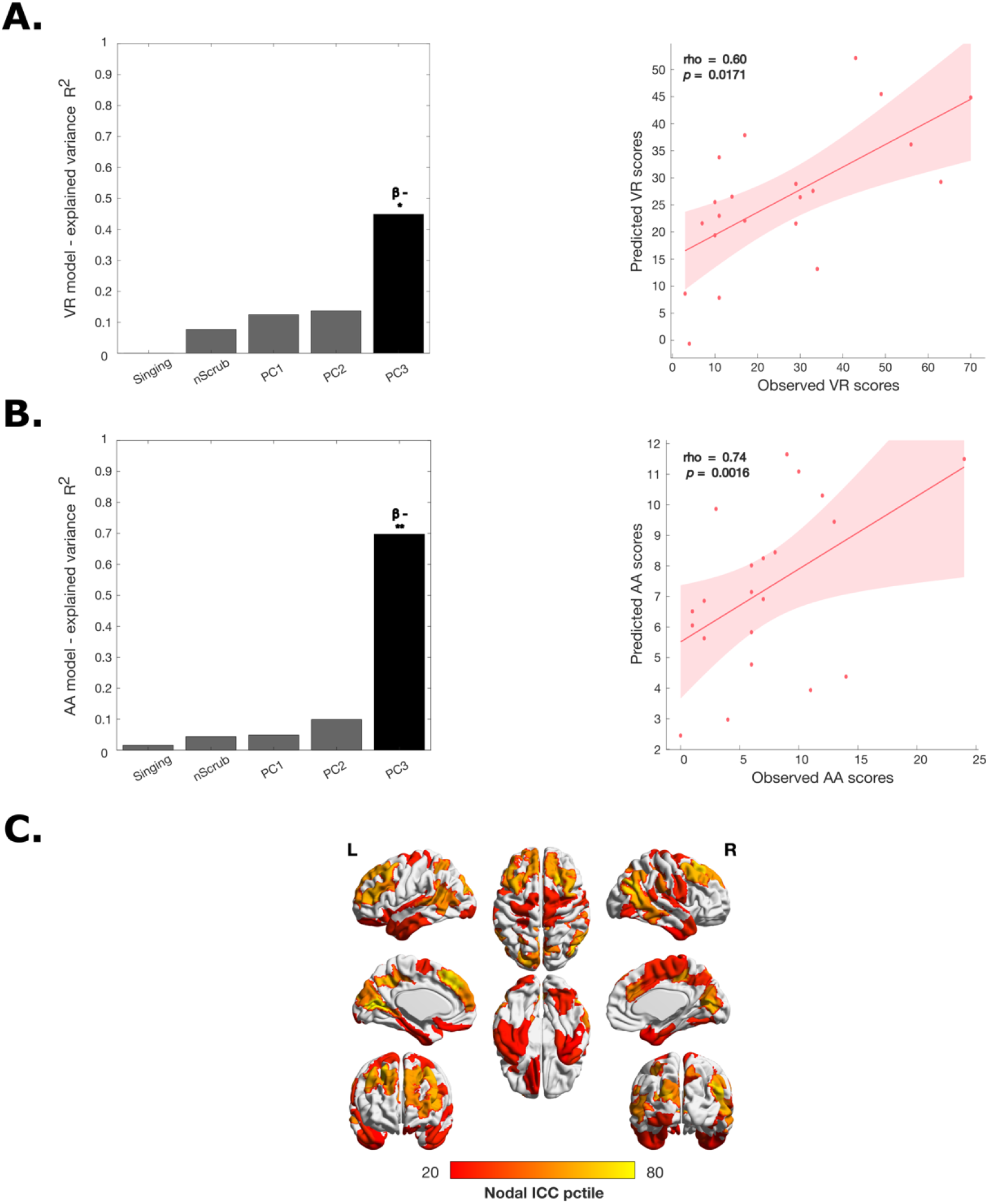
Peak predictive multi-linear model of subjective effects. Following an ICC derived feature selection comprising k-fold validation and null-modelling (see Methods), 3000 edges were found to yield explanatory power. **(A)** Visual restructuralisation (VR) peak performance model. Left: The additive linear model consists of two nuisance variables (nScrub, Singing), and three PCA predictors (PC1-3); Right: Scatter plot of the Observed VR scores versus predicted model VR scores. **(B)** Auditory alteration (AA) peak performance model. As before, left: incorporated predictors in an additive linear model, Right: observed vs predicted AA scores. **(C)** Nodal strength (sum across rows) render of top predictive edges. Percentiles are shown (from 20th to 80th percentile). For all panels, significant predictors are denoted as follows: *p*<0.05*\*, p*<0.01** with β-indicating that its beta coefficient is negative.

### Additional control analyses

For completeness, we performed a series of quality controls on our primary identifiability analysis. More specifically, we (i) repeated our main analyses using censored fMRI timeseries, (ii) assessed differences in motion metrics between conditions, (iii) examined split-half differences in primary motion outcomes per scan, (iv) evaluated their association with all sFC and dFC identifiability outcomes. Our findings appear robust to motion and replicable across different denoising strategies (Figure.S2).

## Discussion

Here, we leveraged the understanding that an individual’s functional brain connectivity profile is both unique and reliable, to document how the inherent features of a subject’s functional connectome might transition into a collective altered state of consciousness. Using the concept of connectome fingerprinting outlined by Amico et al.^11^, in a cohort of 21 Santo Daime members taking part in the religious use of ayahuasca, we were able to detect for each subject a significantly greater proportion of shared functional connectivity traits across different timescales of neural processing. Furthermore, we show that this shared variance is accompanied by the reconfiguration of keypoint edges pertinent to higher-order functional subsystems, otherwise driving normative brain “fingerprints”. Equally, we show that the instability of edges is likely relevant to experiential differences given that they can be used to predict aspects of an individual’s subjective experience. Together, these findings may reflect the presence of a common functional space among churchgoers, in which the general blueprint of a subject’s inherent resting-state functional connectivity shifts to overlap with fellow members (see Fig.7 for an analogy). Ultimately, subject-level approaches such as those presented herein highlight the potential to discover personalised fMRI-based connectivity markers that may eventually be used to trace a subject’s functional connectome across states of consciousness.

**Figure 7.**
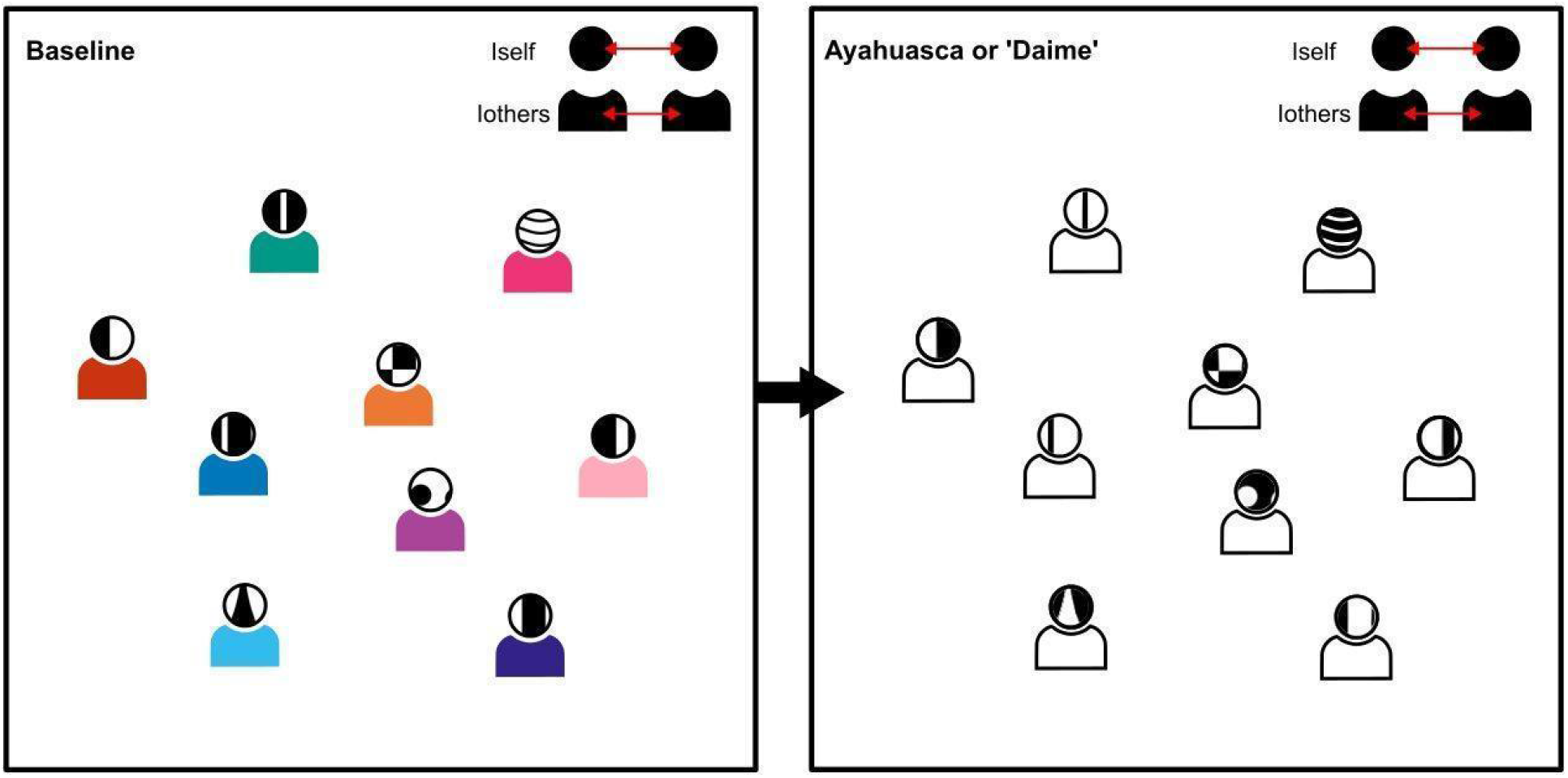
Outfit swap analogy of changes to subject variability under ayahuasca. In everyday life people rarely dress one and the same. Ordinarily, an individual might choose their attire based on personal preference, such as a colourful shirt. The colour palette that we might choose would represent our distinctiveness, in turn differentiating us from others (I_others_). Throughout our day, I_others_ might vary, given others with distinct preferences might come and go. Now say in a different scenario, such as a Santo Daime ceremony, we were to abide by the dress-code of a white uniform. Even if others might come and go, our similarity to others (I_others_) at the ceremony would be high since the colour white is mandated throughout the event for all participants. However, if the uniform were to be contrasted to daily life (I_othersHybrid_) we might find it to be just as dissimilar as any other coloured shirt that we might come across on an average day. In parallel the constituents of our self-identity (I_self_) may equally be denoted as a unique pattern. Whereas their total variability is unlikely to change regardless of circumstance, an acute perturbation by a pharmacological agent such as ayahuasca may lead to its reconstitution (I_selfHybrid_).

### The collective use of ayahuasca yields shared functional traits

Whereas we found the I_diff_ of everyone’s connectome was diminished under ayahuasca, this reduction could be ascribed to the increased contribution of I_others_. In other words, a subject’s brain fingerprint holds a larger number of shared functional traits under ayahuasca, diminishing its overall differentiability. When factoring in the practices of Santo Daime, this could echo the collective “works” carried out by church members. While ayahuasca experiences are highly subjective, followers engage in a set of stereotyped behaviours such as singing when under the influence. Evidence from classical psychedelics suggests an ‘unconstrained’ state of cognition of few deliberate or automatic constraints, featuring a large amount of hyper-associative thinking and diminished reality-testing ^46^. However, contextual components may inherently tether these internal states and with that, their underlying FC profile. Using cognitive strategies such as attentional deployment in order to attend to a religiously meaningful world, members of Santo Daime might instead partake in a constrained state of cognition. For example, prior imaging work with church-goers has previously drawn parallels with the induction of a task-active state, exemplified by suppressions in DMN activity ^47^ normally associated with external goal-directed attention such as task engagement or focused attention meditation ^48,49^. The overlaps between serotonergic psychedelics and meditation are often a recurrent theme ^50^ and when studied in tandem, have been shown to modulate subacute functional coupling of the DMN ^51^.

An interpretation could be that an internal labour bound by a religious framework is disseminated across individual connectivity matrices akin to a common functional space. Tellingly, evidence from studies with normative samples, show that the inter-subject variability of a sample is diminished when engaging in a task battery, proportionally to cognitive load ^14,31^. Given increases in dI_others_ (and accordingly, reductions in dI_diff_) across temporal scales solely manifest beyond 210 seconds, shared functional traits could be interpreted as indeed residing in previously identified timeframes at which complex cognition emerges ^25^. However, without future work disentangling the many spontaneous cognitive processes arising during resting-state from specific patterns or windows of dynamic FC, the significance of this interpretation is uncertain. More so than tasks, “ground truth” approaches for pharmaco-imaging such as films ^52^ or other integrated designs ^53^ paired with subject-level dynamical analysis approaches ^54^ may hold promise in tagging the behavioural relevance of dFCs for attention-impairing drugs such as these.

### Constituents of connectome self-identity are mutable under ayahuasca

Studies have repeatedly demonstrated the remarkable consistency of inherent functional connectivity patterns across participants and mental states ^24,55^. The majority of a subject’s uniqueness or inter-session variance (between 63 and 87%) can be explained by commonalities in functional connectivity architecture between states ^14^. Here, we show that contrary to a phenomenological loss of self-identity, self-identifying connectivity is also preserved under ayahuasca. Psychedelics are described to produce a wide-scale discoordination of brain activity ^56,57^ denoted by a structural-functional uncoupling ^58^. To take the view that fingerprinting comprises fixed anatomical loci which assimilate several information sources to plan coherent behavioural responses ^59^ then their impaired integration as observed under psychedelics should also lead to a diminished I_self_. However, connectome fingerprinting of clinical populations exhibiting structural-functional uncoupling show no differences in I_self_ against healthy controls ^60^. Instead, it is now known the total blueprint of functional connectivity does not constitute discrete networks but is rather best described by more mutable local and global gradients ^61^ which are likely susceptible to pharmacological perturbation. Indeed, our finding of diminished I_selfhybrid_ may instead reflect a general functional reconfiguration of inherent signalling traits, synonymous with the apparition of a novel functional connectivity architecture.

### Local shifts in functional connectivity stability drive altered connectome fingerprints

At a fundamental level, we also looked at the edgewise contributors to identification which might explain the observed reconfiguration of identifiability. Previous work has closely implicated temporal stability of regional functional connectivity as driving connectome fingerprints ^25^. Using ICC, we observed global reductions in edge stability at all measured timescales under ayahuasca, striking proxy of the patterns of functional change under psychedelics. 5-HT2A agonists have been found to produce brainwide increases in signal complexity ^62,63^. which may consequently limit the temporal concordance of edge pairs. Before continuing, it should be noted that functional connectivity and ICC are not interchangeable but rather complementary methods ^64^ and future work should continue to examine the mediating relationship between the differing measures of complexity available, ICC and connectome identifiability ^65^.

Given that certain edges drive a subject’s normative fingerprint, we examined how regional contributions to identifiability evolve under ayahuasca. While a subset of 250 edges could maximally define a subject’s fingerprint, their importance markedly dropped under ayahuasca. Disseminated across higher-order association cortices these regions are shown to encode the majority of inter-individual variance ^12,37^. Importantly, it has been previously hypothesised that the appearance of a desegregated functional architecture under psychedelics stems from the impairment of these same functional subsystems ^35^. However, frontotemporal DMN nodes central to the effects of hallucinogens ^66^ emerged as the focal point for a subject’s identifiability under the influence, expressing greater stability. The DMN has been implicated in different aspects of conscious experience, such as ongoing cognition ^67^, spontaneous thought ^68^, rumination ^69^, and self-referential processing ^70^ and it’s select prevalence may further highlight a diminished variability of subjective experience ^71^.

These shifts in stability were also pronounced at a network level. While functional networks do not equally contribute to an individual fingerprint, each functional subsystem is thought to have temporal “peaks’’ in stability ^25,37^. Here, links overall pertaining to VIS and SM between-network connectivity exhibited greater desynchrony under ayahuasca, varying across all examined temporal windows. These findings appear in line with patterns of hyper-connectivity in sensory networks that have been observed under psychedelics ^72,73^, thought to reflect a de-differentiation of hierarchical organisation ^74^. Furthermore, clustering-based dFC approaches with LSD and psilocybin have shown an increased fractional occurrence and dwell time of alternating states of hyperconnectivity ^33,75,76^ which may account for the stochasticity of edge stability at each temporal window. It could be therefore suggested that the outcome of a dedifferentiation of functional hierarchies under psychedelics may also extend to their temporal organisation. Suggesting this, both ends of the continuum of hierarchical organisation became temporally more similar to one another under ayahuasca; with unimodal regions (ie: parietal operculum, visual cortex) associated with rapid multisensory processing now peaking in stability at longer timescales and transmodal regions (ie: prefrontal cortex, posterior cingulate cortex) otherwise exhibiting longer, integrative firing patterns maximal at shorter timescales. Applying approaches to examine the temporal propagation and latency of brain activity ^77,78^ under psychedelics may highlight new explanations for their effects on network architecture.

### Connectome fingerprints are relevant to the subjective ayahuasca experience

Assuming ayahuasca experiences are highly individual, might subject-level shifts in functional connectivity also help predict overlaying subjective experiences? To explore this hypothesis, we devised a data driven PCA approach to assess the behavioural relevance of highly identifiable edges. Our results suggest that subsets of highly stable edges not only drive a subject’s identifiability under the influence but also hold explanatory power for the AUD and VIS dimensions of the 5D-ASC. In practice, by decomposing the total variance of a FC signal to a reduced number of orthogonal components, PCA offers the opportunity to separate the contribution of subject-level and group-level functional connectivity information carried in low and high-variance components. If group-differences were highly explanatory of experiential scores, then a single, highly explanatory factor (PC1) might have emerged as a principal model contributor. Instead, our results were contingent on the inclusion of PC3 as a predictor of interest, suggesting higher-order PC deviations capturing individual variation in FC were most relevant to the visual and auditory effects of ayahuasca. Furthermore, predictive edges were found to span primarily both higher order systems (DMN, FPN) and primary systems (VIS, SM), with the former contributing more (in number) to behavioural prediction as per prior work ^12,21,37^. Given their developmentally late maturation ^79^, susceptibility to individual environmental effects ^80^ and, dense 5-HT2A expression ^81^ and coordination of multisensory integration in comparison to primary systems ^61^, higher-order regions may more easily account for divergent phenomena, more so than primary systems, themselves partially influenced by the temporary states of each individual during scanning ^82^.

## Limitations

The present work comes with several limitations. Importantly, members of Santo Daime are not reflective of the general population. Drinking ayahuasca several times a month, members likely exhibit a level of habituation to the drug’s effect. Furthermore, 5-HT_2A_ agonists are potent psychoplastogens ^83^ likely inducing structural alterations after prolonged use. For example, cortical thickness analyses of Santo Daime members have demonstrated an association between significant thinning in midline structures and self-transcendent personality traits ^84^. There is an abundance of studies showing how between-subject differences in white matter integrity are intimately related to interindividual variability in functional dynamics ^85,86^ and likely identifiability ^87^. As with observational studies, these findings are subject to confounding effects regarding dosage, blinding, sample inclusion criteria and expectancy. Adequate blinding in studies comprising experienced users continues to be an unresolved factor in the field, due to a subject’s immediate recognition of a drug’s effect (or non-effect). This is further accentuated here, given the ritual elements. Future studies with a larger diverse sample of ceremonial users may benefit from not only counterbalancing and adequately blinding the agent in question, but also surrounding practices mediating experiential outcomes. With regard to the methodology, it is well-known that head motion due to its potential for skewing functional connectivity estimates ^88,89^ is likely a confounder in the study of inter-individual differences ^12^. If treated as a “state” characteristic for subjects, joint differences among members of a group might account for a proportion of between-subject variability. Whereas numerous steps were taken to exclude its influence, it is unknown to what degree factors such as motion, respiratory fluctuations or arousal level may prevail in shorter dFC windows. Future studies should aim to replicate this workflow using framewise approaches such as dynamic conditional correlations ^90^ or phase coherence estimation ^91^. Per prior work ^25^, preprocessing was performed with a pipeline comprising Global Signal regression (GSR). GS is hypothesised to contain a complex mixture of non-neuronal artefacts (e.g., physiological, movement, scanner-related) and its removal, while effective, is widely debated In light of differing results for psychedelic effects on resting-state measures ^47,56,73^ following its use, a consensus on the suitability of GSR alongside other preprocessing steps for pharmacological neuroimaging should be reached ^92^.

## Conclusion

In summary, the ritualistic use of ayahuasca produces a shared functional space, marked by a spatiotemporal reconfiguration of brain connectivity traits. Members of Santo Daime pertain to a culture which emphasises the interaction of a psychoactive sacrament with the interpersonal dynamics of ritualism. Ultimately, it is likely the synergy of the two that produces the blurred connectome fingerprint presented herein. An important next step is to replicate the presented approach to double-blind imaging data with other classical psychedelics. It may very well be the case that for normative samples, the opposite holds true, with interindividual differences being accentuated as a result of a diversified content of thought. Going ahead, by celebrating individual differences in the study of subjective experiences we may be a step closer to producing personalised neural markers of psychedelic effects.

## Methods

### Participants

Twenty-four volunteers were enrolled in a within-subject, fixed-order observational study. Data from three volunteers were excluded from analyses due to excessive head motion leaving a final sample of 21 subjects (10 females) of ages 29 to 64 (*M*: 54.48, *SD*: 10.55). The cohort consisted of experienced members of the Dutch chapter of the church of Santo Daime. Individuals were selected based on an exclusion criterion comprising absence of ferromagnetic devices/implants (MRI contraindications), pregnancy and use of (medicinal) substances in the past 24 h. Detailed demographic information pertaining to the final sample can be found in Table S1.

All participants were fully informed of all procedures, possible adverse reactions, legal rights and responsibilities, expected benefits, and their right to voluntary termination without consequences.The study was conducted according to the Declaration of Helsinki (1964) and amended in Fortaleza (Brazil, October 2013) and in accordance with the Medical Research Involving Human Subjects Act (WMO) and was approved by the Academic Hospital and University’s Medical Ethics committee (NL70901.068.19/METC19.050).

### Study procedures

Participants underwent two consecutive test days; one baseline (sober) followed by one under the influence of ayahuasca.

Participants self-administered a volume of ayahuasca equivalent to their usual dose (mean 24ml, *SD*: 8.16), prepared from a single batch by the Church of Santo Daime and analysed according to prior referencing standards (see Supplementary). As to facilitate the communal use of ayahuasca, participants drank ayahuasca brew while initiating the works in company of fellow members. Participant dosing schedules were stratified across each lab visit with testing performed within 4 pairs of visits (6 subjects per cycle) with each subject being tested at the same window of time as to minimise diurnal variation. The brew used contained 0.14 mg/ml of DMT, 4.50 mg/ml of harmine, 0.51 mg/ml of harmaline, and 2.10 mg/ml of tetrahydroharmine. Each ceremony was organised and supervised by the Santo Daime church. The research team was not involved in the organisation of the ceremonies nor the production, dosing, or administration of ayahuasca.

On each day upon arrival to the lab, absence of drug and alcohol use was assessed via a urine drug screen and a breath alcohol test. An additional pregnancy test was given if participants were female. Each visit consisted of a 30-minute wait period, followed by a 1h MRI scanning session occurring 1h after intake. On day 2, venous blood samples were collected approximately 60 and 160 minutes after ayahuasca intake to assess serum concentrations of alkaloids according to laboratory protocols (Supplementary). The retrospective 5-Dimensions of Altered States of Consciousness (5D-ASC) scale ^93^ and the Ego Dissolution Inventory ^94^ were administered 360 minutes after drug ingestion to assess the subjective experience after drug intake (for details, see Supplementary). Each visit to the lab lasted 6 hours.

Following study completion, each subject was contacted for an online follow-up. In order to gauge the prevalence of mental processes pertaining to the ceremonial use of Daime during resting-state participants were asked to answer visual analogue scales (0-100) pertaining to their *recollection* of each resting-state acquisition, whether they were internally *singing* during, or employing *meditation* in the scanner. For more information regarding all procedures, inventories, and corresponding subscales, please see the Supplementary.

### Image acquisition

Images were acquired on a MAGNETOM 7T MR scanner. On each visit, participants underwent a structural MRI (60 min post treatment), single-voxel proton MRS in the PCC (70 min post) and visual cortex (80 min post), and fMRI (90 min post), during peak subjective effects. Findings and methods pertaining to MRS are to be reported elsewhere.

T1-weighted anatomical images were acquired using a magnetisation-prepared 2 rapid acquisition gradient-echo (MP2RAGE) sequence (TR = 4500 ms, TE = 2.39 ms, TI1 = 0.90 s, TI2 = 2.75 s, flip angle 1 = 5°, flip angle 2 = 3°, voxel size = 0.9 mm isotropic, matrix size = 256 × 256 × 192, phase partial Fourier = 6/8, GRAPPA = 3 with 24 reference lines, bandwidth = 250 Hz/pixel). 500 whole brain echo planar imaging (EPI) volumes were acquired at rest (TR = 1400 ms; TE = 21 ms; field of view=198 mm; flip angle = 60°; oblique acquisition orientation; interleaved slice acquisition; 72 slices; slice thickness = 1.5 mm; voxel size = 1.5 × 1.5 × 1.5 mm) followed by 5 phase encoding volumes acquired for EPI unwarping. During EPI acquisition, participants were shown a black fixation cross on a white background.

### Functional preprocessing

All preprocessing steps were performed according to an in-house pipeline ^95,96^ based on FSL (FMRIB software library, FSL 6.0; www.fmrib.ox.ac.uk/fsl) and implemented in MATLAB (R2019b). The individual functional connectomes (FCs) were modelled in the native BOLD fMRI space of each subject.

MP2RAGE images were first denoised to improve signal-to-noise ratio ^97^, bias-field corrected (FSL FAST), skullstripped (HD-BET)^98^, and then segmented (FSL FAST) to extract white matter, grey matter and cerebrospinal fluid (CSF) tissue masks. These masks were warped in each individual subject’s functional space by means of subsequent linear and non-linear registrations (FSL *flirt* 6dof, FSL *flirt* 12dof and FSL *fnirt*). BOLD fMRI volumes were preprocessed in line with Power at al.^88,89^. Subsequent steps included: deletion of 2 initial volumes (FSL utils), slice timing correction (FSL slicetimer), BOLD volume unwarping (FSL topup), realignment (FSL mcflirt), normalization to mode 1000, demeaning and linear detrending (Matlab detrend), regression (Matlab regress) of 18 signals: 3 translations, 3 rotations, and 3 tissue-based regressors (mean signal of wholebrain, white matter (WM) and cerebrospinal fluid (CSF)), as well as 9 corresponding derivatives (backwards difference; Matlab). A bandpass first-order Butterworth filter [0.009 Hz, 0.08 Hz] was applied to all BOLD timeseries at the voxel level (Matlab butter and filtfilt). As a final denoising step, the first three principal components of the BOLD signal in the WM and CSF tissue were regressed out of the gray matter (GM) signal (Matlab, pca and regress) at the voxel level. No smoothing was performed. We also kept track of the fMRI volumes that were highly influenced by head motion, by using three different metrics as a scrubbing index: 1) Frame Displacement (FD, in mm); 2) DVARS (D referring to temporal derivative of BOLD time courses, VARS referring to root mean square variance over voxels)^88^;3) SD (standard deviation of the BOLD signal within brain voxels at every time-point). The FD and DVARS vectors (obtained with fsl_motion outliers) were used to detect outlier BOLD volumes with FD > 0.3 mm and standardized DVARS > 1.7. The SD vector calculated using Matlab was used to detect outlier BOLD volumes higher than 75 percentile + 1.5 of the interquartile range per FSL recommendation^99^. It should be noted no volume censoring was performed using this index. Rather, this information was used as a confound in our multilinear regression analyses and quality control assessments (see Fig.5 and Fig. S2). Functional connectomes obtained with and without scrubbing were highly similar (average Pearson’s r = 0.99) with no significant differences in motion being identified between or within conditions (see Fig. S2).

A 2mm cortical Schaefer parcellation^100^ based on 200 brain regions (publicly available at: https://github.com/ThomasYeoLab/CBIG/tree/master/stable_projects/brain_parcellation/Schaefer2018, was projected into each subject’s T1 space (FSL flirt 6dof and FSL flirt 12dof) and subsequently their native EPI space. FSL boundary-based-registration was also applied to improve the registration of the structural masks and parcellation to the functional volumes. Regions of interest (ROIs) were ordered according to seven cortical RSNs as proposed by Yeo et al.^157^. These included the visual (VIS), somatomotor (SM), dorsal attention (DA), ventral attention (VA), limbic (L), frontoparietal (FP) and the default mode network (DMN).

### Assessment of functional connectivity

In order to assess connectome fingerprints (described in our next section) across static and dynamic temporal scales, we devised two separate workflows for functional connectivity. Static functional connectivity between each pair of ROIs was calculated with a Pearson correlation coefficient (MATLAB corr) between each pair of mean signal time courses (across a single run). For each subject, this results in an *N* × *N* FC matrix, where N is the number of ROIs, and with each element in the FC representing the connectivity strength between a pair of ROIs. Secondly, we assessed the dynamic functional connectomes (dFC) by performing a sliding window analysis to produce sets of connectivity matrices reflecting the temporal development of whole-brain functional connectivity (across our 249 timepoints)^25^. We captured relevant FC patterns by balancing the number of time points for a stable dFC computation, exploring sets of dFCs across 5 different window lengths of: 70s, 140s, 210s, 280s and 349s. Each window step was fixed to 14s, the equivalent of 10 fMRI data points.

### Whole-brain connectome identifiability

Changes to the identifiability of each subject’s functional connectome were quantified by replicating the methodology originally proposed by Amico et al., devised for both static and dynamic functional connectivity ^11,25^.

The approach devises an identifiability matrix for each condition, consisting of a matrix of correlations (Pearson, square, non-symmetric) between a subject’s test and retest functional scans. We firstly split each scan into two corresponding halves (249 volumes each, or 6 min) to generate test-retest sets for each condition. Prior work has shown fMRI scan lengths of 3 minutes are sufficient to produce reliable fingerprints ^11,20^. Since connectivity matrices are symmetric, we can then extract unique elements of each test-retest FC by taking the upper triangle of each matrix; resulting in a 1×19900 vector of edge values for each subject per condition which can then be compared using Pearson correlation, either between different subjects in the same condition or within the same subject across conditions. This yields the “identifiability matrix” as outlined in Figure 1.

In the case of static FCs, let **A** be the “identifiability matrix”, between the subjects’ FCs test and retest. The dimension of **A** is N^2^, where N is the number of subjects in the study. Let I_self_ = <a_ii_> represent the average of the main diagonal elements of **A**, which consist of the Pearson correlation values between test-retest sets of same subjects, otherwise defined as self-identifiability or I_self_. Similarly, let I_others_ = <a_ij_> define the average of the off-diagonal elements of matrix **A**, i.e. the correlation between test-retest sets of different subjects. Lastly, let the differential identifiability (I_diff_) of the population be the difference between both terms, otherwise denoted as:

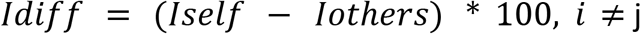

Which provides an indication of the difference between the average within-subject FCs similarity and the average between-subjects FCs similarity. The greater the I_diff_, the higher the differential identifiability overall along the population. As an additional step, we also sought to derive distance of each participant fingerprint under Ayahuasca (ie. I_self_, I_others_ I_diff_) from their respective normative state from their baseline. Using the approach outlined by Sorrentino et al for static connectomes^44^ we calculated the identifiability matrices across combinations of different conditions (the Pearson correlation of test-sober, retest-ayahuasca). When concatenated with our within-group identifiability matrices this produces a hybrid identifiability matrix (see figures 1 and 2), where the between blocks (groups) elements and scores reflect the similarity (or distance) between the test-retest connectomes of subjects across different conditions. By averaging, this also allows us to derive a final overall cohort I_clinical_ score which provides a percentage (average) score of how similar their connectome with respect to baseline is. Finally, we also measured the Success-rate (SR) of the identification procedure as percentage of cases with higher I_self_ vs I_others_ ^12,101^. For completeness, we calculated per condition the significance of both observed Idiff and SR scores in respect to their null equivalents using permutation testing (see Supplementary).

We can next extend this principle to dynamic functional connectomes (dFC) by calculating each measure across each dynamic frame of connectivity (see figure 1 for an overview). For a fixed window length *w*, the resulting dynamic identifiability matrix is then a block diagonal matrix, where each block represents the self-similarity within the dFC frames of a specific subject. The off-diagonal blocks, in this representation, encode instead the between-dFC frames similarity across different subjects (dynamic I _others_). Let 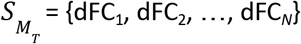 be the set of dFC frames in the test session for a specific subject *M*. Similrly, Let 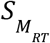 represent the set of dFC frames in the retest session for the same subject *M*. We can then define the dynamic Iself (dI_self_) for subject *M* as:

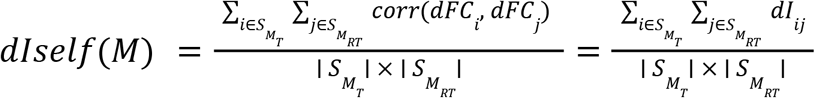

Whereby 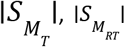 define the cardinalities of the sets. Similarly, let 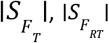 define the sets for a different subject *F*. We can define dynamic Iothers (dI_others_) as:

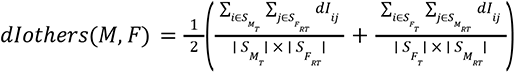

and hence:

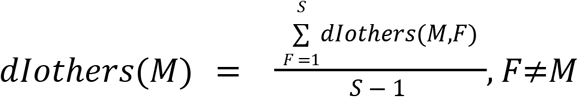

where the summation is over the total number of subjects *S* other than *M*. Last, dI_diff_ for a subject *M* can be described as:

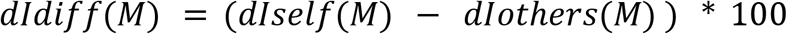

### Edgewise connectome identifiability

In order to understand which edges were key contributors to changes in connectome identifiability, we quantified the edgewise reliability of individual connectomes using intraclass correlation analysis as per^11^. Coefficients derived from ICC are widely used as a reliability index in test-retest analyses, reflecting the percentage agreement between two units of measurement within the same group^102^. The greater the ICC value, the greater the consistency these two units hold. For reference, ICC values below 0.40 are suggested to be poor/unreliable whereas those beyond 0.90: excellent/congruent^103^.

We employed this approach under the assumption that subsets of highly synchronous, or stable edges across test-retest sets edges are major drivers of each state’s connectome fingerprint. For a FC, this generates a square symmetrical ICC matrix of size N^2^, where N is the number of brain regions (see figures 2 and 4) for a specific timeframe. From this, we can also extrapolate network identifiability by averaging ICC values of within and between network edges, producing 7 × 7 ICC fingerprint matrices corresponding to our Yeo parcellation. Note that edges were thresholded according to lower bounds of ICC (0.40).

In the case of dynamics, there might be FC frames where identification is higher than others. Consequently, this might not reflect the average behaviour depicted by dI_diff_ thereby skewing ICC estimates. To cover the necessity of that, for each subject session, we sorted the dFC frames in test-retest according to their similarity, from highest to lowest, based on their dIself_j_ value (see *d*Iself equation above). We then recalculated dI_self_, dI_others_, and dI_diff_ when iteratively adding dFC frames one at the time, starting from the best matching ones and then proceeding based on their similarity values in order to end with a “top” frame for each timescale on which our ICC analyses could be performed. As a supplementary analysis we also examined the relationship between FC variability and stability by calculating the standard deviation of functional connectivity for each frame (see figure S4.).

With the expectation that subsets of static edges might primarily contribute to each condition’s identifiability, we sorted these according to their thresholded ICC values computed on the sober condition (baseline). Edges were added in a descending fashion, with I_diff_ being recalculated at each iteration of 50 edges. We selected the sober condition as an index in order to visualise the evolution of normative drivers maximally contributing to Idiff.

### Edgewise prediction of subjective experience

In light of the individual nature of subjective experience and connectome fingerprints we opted for an iterative multilinear modelling approach (MLR) similar to connectome predictive modelling^3^, which instead of summarising predictive edges, uses a principal component analysis (PCA)^104^ of highly identifiable edges.

PCA is an unsupervised exploratory approach which is typically used for dimensionality reduction and pattern recognition. By geometrically projecting FCs onto a two-dimensional space, the total variability in individual participant functional connectivity can be placed systematically along one or more of the principal axes as an emergent set of principal components (PCs) ranked according to variance. If a group is relatively homogenous then PCA only generates one PC along which all participants can be mapped. However, if there are systematic differences within the cohort, then one or more statistically independent factors accounting for subject-level information emerges. This decomposition was applied in an iterative fashion for model selection. Firstly, all edges pertaining to the Ayahuasca condition were sorted in descending order according to their ICC value. At each iteration of 50 edges, we performed a PCA decomposition of their functional connectivity values across subjects, retaining three PCA components ranked according to explained variance. Then, a MLR was built for each subjective effect measure, comprising these PCA components as predictors of interest alongside two covariates: singing (self-reported internal singing during the resting-state) and scrubbing (number of valid volumes). Absence of multicollinearity was assessed using variance inflation factor (VIF)^105^. At each iteration, we strengthened the reliability of our model using the k-fold cross-validation^106^ with k= 5. Specifically, k iterations were performed and at each iteration the k^th^ subgroup was used as a test set. For each iteration, the Spearman’s correlation coefficient between predicted and actual inventory values was calculated and considered as a performance score. We assessed the reliability of this performance score against surrogate models, computed using a set of randomly permuted edges at each step. For each variable of interest this process was repeated 100 times. In order to further reduce the risk of overfitting, modelling was only performed on static edges.

### Statistics

Statistical analyses were carried out in MATLAB 2019b. Shapiro-Wilks was firstly used to assess the normality of all measures. Control variables (subjective effects, PK) were assessed by means of one-tailed t-tests against zero. All outcome measures were analysed in a two-tailed fashion according to their normality; either by Wilcoxon sign-rank (*W)* or Student t-tests (*t)*. Observed static identifiability scores (true) values were examined against corresponding null-distributions following a permutation testing framework (see supplementary). Regarding network-based statistics, we retrieved a Bonferroni-corrected p-value according to the number of unique elements in each matrix. The alpha criterion of significance for all inferences was set at p<0.05.

## Data availability

The connectomes and the accompanying covariates used to differentiate individuals can be made available to qualified research institutions upon reasonable request to J.G.R and a data use agreement executed with Maastricht University.

## Code availability

All code used for analysis is to be made available on P.Mallaroni’s GitHub page (https://github.com/PabloMallaroni/Ayahuasca_Fingerprints).

## Author contributions

J.G.R, K.O and N.M conceptualised the study. L.K, N.M, J.T.R and P.M collected the data. S.T performed all pharmacokinetic analyses. P.M performed all functional connectivity analyses. E.A. devised the identification framework and contributed data analysis tools. H.T. and P.M devised the model analyses. E.A, J.G.R and N.M and provided guidance with methods and data interpretation. P.M wrote the first draft of the manuscript. All authors contributed to the editing of the manuscript.

## Acknowledgments

We would like to thank Gregory Cooper and Emanhuel Troizi Lopez for insightful conversations on the nature of our findings. We are grateful for the extended cooperation of members of the church of Santo Daime.

