## Supplementary for "Ritualistic use of ayahuasca enhances a shared functional connectome identity with others"

**Subjective effect scale descriptions.**
*5D-ASC*. The 5-Dimensions of altered states of consciousness questionnaire contains 94 items in the form of 0-100 visual analogue scales (VAS). The instrument consists of five dimensions comprising Oceanic Boundlessness, Anxious Ego Dissolution, Visionary Restructuralisation, Vigilance Reducton, Auditory Alterations and 11 subscales (Studerus, Gamma et al. 2010). The dimension "Oceanic Boundlessness" (27 items) comprises feelings of derealisation and depersonalization associated with positive emotional experiences. Its lower order scales consist of "experience of unity," "spiritual experience," "blissful state," and "insightfulness." The dimension "Anxious Ego Dissolution" (21 items) feeling of ego-loss and self-control phenomena associated with anxiety. Its lower-order scales are "disembodiment," "impaired control of cognition," and "anxiety." "Visionary Restructuralisation" (18 items) describes perceptual changes, consisting of the subscales "complex imagery," "elementary imagery," "audio-visual synesthesia," and "changed meaning of percepts." Lastly, "Auditory Alterations" and "Reduction of Vigilance" consist of 15 items and 12 items respectively.

*EDI.* The Ego Dissolution Inventory is an eight-item, self-report 0-100 VAS scale that assesses the participant's experience of ego dissolution (Nour, Evans et al. 2016). Sample items for the scale include the following: "I experienced a dissolution of my 'self' or ego" and "I felt at one with the universe." The higher the total score, the stronger the experience of ego dissolution.

**Retrospective resting-state mentation.**
Following study completion, participants were asked to characterise their inner experience during each resting state (at baseline and acutely). Three 0-100 VAS scales were delivered by Qualtrics XM12, with participants asked to specifically recollect their time *looking at the black cross on the white screen* (ie. the resting-state acquisition). These comprised questions pertaining to activities relevant to the *"works"* that Santo Daime members undertake during ceremonial intake (Hartogsohn 2021) as well as their recollection confidence of each session:1) *"How confidently do you remember your brain MRI scanning sessions with us?,* 2)*"When looking at the cross in the MRI scanner, did you think of Daime hymns or sing them in your head?",* 3)"*When looking at the cross in the MRI scanner, did you purposely try and enter a state of meditation? For example, focusing your mind to quieten your thoughts?".*

**Ayahuasca brew alkaloids**
A sample of the ayahuasca batch used for each ceremony was collection.The alkaloid concentrations of the brew were determined as in previous studies (Kiraga, Mason et al. 2021) after dilution and extraction (1-chlorobutane/ether, 1:1, v/v) utilizing LC–MS/MS (Agilent, Waldbronn, Germany), calibrated with pure reference substances of DMT (Cerilliant, Round Rock TX, US), harmine, harmaline and tetrahydroharmine (Aldrich Chemistry, St. Louis MO, US).

**Pharmacokinetic measures**
Venous blood samples were collected at 60 and 160 minutes after ayahuasca ingestion. Samples were aliquoted, encoded, and frozen at −80 °C until they were used. Analysis of the alkaloids DMT, harmine, harmaline and tetrahydroharmine in serum samples (200 µl) was performed after extraction with ethyl acetate using LC–MS/MS (Agilent, Waldbronn, Germany). Calibration curves covered the range 0.25 – 40 ng/ml with lower limits of quantitation (LLOQ) of DMT 0.077, harmine 0.13, harmaline 0.23, and tetrahydroharmine 0,18 ng/ml.

**Table S1.** Demographics characteristics and ayahuasca use history (N = 21).

| Variable | M ± SD |
| --- | --- |
| Gender (male/female) | 11/10 |
| Age, years | 54.48 ± 10.55 |
| Body mass index (kg/m$^2$) | 23.54 ± 3.47 |
| Member of Santo Daime, years | 14.52 ± 8.66 |
| Lifetime ayahuasca use, number of occasions | 562.62 ± 683.73 |
| Time since last ayahuasca use, days | 54 ± 101.78 |
| Dose, ml | 24 ± 8.16 |

Imaging sample demographics. Means (M) and standard deviations (SD) are listed where applicable.

**Table S2.** Subjective effect markers of ayahuasca experience. (N = 21).

| Inventory | M(SD) | SEM | t(20) | Cohens *d* | $P_{zero}$ |
|---|---|---|---|---|---|
| *5D-ASC (% score)* | | | | | |
| Oceanic boundlessness | 33.1±20.79 | 4.54 | 7.3 | 1.59 | **<0.0001** |
| Anxious ego-dissolution | 5.1±4.78 | 1.04 | 4.88 | 1.06 | **<0.0001** |
| Visionary restructuralisation | 26.24±20.04 | 4.37 | 6 | 1.31 | **<0.0001** |
| Auditory alterations | 7.24±5.6 | 1.22 | 5.92 | 1.29 | **<0.0001** |
| Reductions of vigilance | 16.24±10.93 | 2.38 | 6.81 | 1.49 | **<0.0001** |
| Experience of unity | 32.67±27.26 | 5.95 | 5.49 | 1.2 | **<0.0001** |
| Spiritual experience | 38.24±26.93 | 5.88 | 6.51 | 1.42 | **<0.0001** |
| Blissful state | 45.29±28.78 | 6.28 | 7.21 | 1.57 | **<0.0001** |
| Insightfulness | 34.9±23.69 | 5.17 | 6.75 | 1.47 | **<0.0001** |
| Disembodiment | 13.19±13.79 | 3.01 | 4.38 | 0.96 | **0.0001** |
| Impaired control and cognition | 7.57±8.32 | 1.82 | 4.17 | 0.91 | **0.0002** |
| Anxiety | 3.09±3.25 | 0.71 | 4.36 | 0.95 | **0.0001** |
| Complex imagery | 31.48±27.29 | 5.96 | 5.28 | 1.15 | **<0.0001** |
| Elementary imagery | 38.9±29.91 | 6.53 | 5.96 | 1.3 | **<0.0001** |
| Audio-visual synsthaesia | 29.38±25.73 | 5.62 | 5.23 | 1.14 | **<0.0001** |
| Changed meaning of percepts | 16.76±19.68 | 4.29 | 3.9 | 0.85 | **0.0004** |
| *EDI (% score)* | | | | | |
| Total | 35.8±22.95 | 5.01 | 7.15 | 1.56 | **<0.0001** |

Means (M), standard deviations (SD) and standard errors (SEM) are listed where applicable. Statistical values for one-tail t-tests against zero for each subscale are presented. Scores are listed as percentage of maximum possible (POMP). Significant differences are listed in **bold.**

**Table S3.** Time course of ayahuasca metabolites in serum (ng/ml), as determined by liquid chromatography-mass spectrometry (LC-MSMS).

| Timepoint | DMT | Harmaline | Harmine | Tetrahydroharmine |
|---|---|---|---|---|
| *+60 min (N=18)* | | | | |
| M(SD) | 18.36±16.17 | 1.57±1.43 | 7.45±7.32 | 21.65±17.23 |
| SEM | 3.53 | 0.31 | 1.6 | 3.76 |
| t(17) | 4.82 | 4.67 | 4.32 | 5.33 |
| Cohens *d* | 1.14 | 1.1 | 1.02 | 1.26 |
| $P_{zero}$ | **<0.0001** | **0.0001** | **0.0002** | **<0.0001** |
| *+160 min (N=19)* | | | | |
| M(SD) | 7.3±6.17 | 1.95±1.52 | 6.85±6.63 | 48.5±26.62 |
| SEM | 1.35 | 0.33 | 1.45 | 5.81 |
| t(18) | 5.16 | 5.59 | 4.51 | 7.94 |
| Cohens *d* | 1.18 | 1.28 | 1.03 | 1.82 |
| $P_{zero}$ | **<0.0001** | **<0.0001** | **0.0001** | **<0.0001** |

Means (M), standard deviations (SD) and standard errors (SEM) are listed where applicable. Statistical values for one-tail t-tests against zero for each timepoint are presented. Significant differences are listed in **bold.**

**Table S4.** Scores of retrospective resting-state mentation items**.**

| | Recollection | Internal singing | Meditation |
|---|---|---|---|
| *TD1 (N=15)* | | | |
| M(SD) | 63.47±25.9 | 24.93±29.14 | 42.87±34.34 |
| SEM | 6.67 | 7.52 | 8.87 |
| *TD2 (N=11)* | | | |
| M(SD) | 66±26.15 | 42.93±36.25 | 37±37.34 |
| SEM | 7.88 | 10.93 | 11.26 |
| *W* | 14 | 58 | 24 |
| *Z* | 0 | 2.25 | -0.8 |
| Cohens *d* | 0.11 | 0.63 | 0.17 |
| *p* | 1 | **0.0261** | 0.4227 |

Means (M), standard deviations (SD) and standard errors (SEM) are listed where applicable. Statistical values for two-tail sign-rank tests are presented. Significant differences are listed in **bold.**

**Table S5.1** Wilcoxon sign rank differences for all network pairs: 70s.

| Edge pair | *W* | *Z* | Cohen's *d* | *p* | Direction |
|---|---|---|---|---|---|
| VIS-VIS | 22165 | -1.04 | 0.05 | 8.3433 | - |
| SM-SM | 36161 | 0.9 | 0.03 | 10.375 | + |
| DA-DA | 8250 | -4.39 | 0.25 | **0.0003** | - |
| VA-VA | 4888 | -0.52 | 0.04 | 16.8088 | - |
| L-L | 180 | 0.47 | 0.05 | 17.8565 | + |
| FP-FP | 17291 | -2.49 | 0.12 | 0.3615 | - |
| DMN-DMN | 71244 | -3.91 | 0.13 | **0.0026** | - |
| VIS-SM | 82429 | -13.64 | 0.47 | **<0.0001** | - |
| VIS-DA | 36882 | -3.16 | 0.12 | **0.0438** | - |
| VIS-VA | 20667 | -7.78 | 0.32 | **<0.0001** | - |
| VIS-L | 699 | -7.02 | 0.42 | **<0.0001** | - |
| VIS-FP | 66388 | -3.3 | 0.13 | **0.027** | - |
| VIS-DMN | 111741 | -1.92 | 0.05 | 1.5344 | - |
| SM-DA | 53697 | -5.57 | 0.18 | **<0.0001** | - |
| SM-VA | 46207 | -0.46 | 0.02 | 18.0656 | - |
| SM-L | 1937 | -3.08 | 0.16 | 0.0573 | - |
| SM-FP | 65582 | -5.78 | 0.17 | **<0.0001** | - |
| SM-DMN | 130698 | -5.84 | 0.15 | **<0.0001** | - |
| DA-VA | 27596 | -2.07 | 0.09 | 1.0853 | - |
| DA-L | 1721 | -2.5 | 0.13 | 0.3512 | - |
| DA-FP | 41670 | -5.28 | 0.19 | **<0.0001** | - |
| DA-DMN | 66604 | -9.09 | 0.27 | **<0.0001** | - |
| VA-L | 1111 | -1.82 | 0.1 | 1.9149 | - |
| VA-FP | 29619 | -5.14 | 0.21 | **<0.0001** | - |
| VA-DMN | 92255 | -3.78 | 0.12 | **0.0043** | - |
| L-FP | 2955 | -3.26 | 0.19 | **0.0315** | - |
| L-DMN | 5245 | -1.18 | 0.06 | 6.7111 | - |
| FP-DMN | 132828 | -5.08 | 0.13 | **<0.0001** | - |

**Table S5.2** Wilcoxon sign rank differences for all network pairs: 140s.

| Edge pair | *W* | *Z* | Cohen's *d* | *p* | Direction |
|---|---|---|---|---|---|
| VIS-VIS | 31638 | 0.02 | 0 | 27.5036 | + |
| SM-SM | 66434 | 2.95 | 0.12 | 0.0878 | + |
| DA-DA | 20747 | 0.58 | 0.03 | 15.7267 | + |
| VA-VA | 6741 | -1.55 | 0.1 | 3.4138 | - |
| L-L | 534 | 0.46 | 0.05 | 18.1723 | + |
| FP-FP | 34183 | 1.14 | 0.06 | 7.0781 | + |
| DMN-DMN | 154675 | -4.13 | 0.12 | **0.001** | - |
| VIS-SM | 161717 | -4.51 | 0.14 | **0.0002** | - |
| VIS-DA | 73504 | -0.31 | 0.01 | 21.0786 | - |
| VIS-VA | 23431 | -10.37 | 0.46 | **<0.0001** | - |
| VIS-L | 6337 | -2.26 | 0.11 | 0.6755 | - |
| VIS-FP | 75226 | -4.6 | 0.16 | **0.0001** | - |
| VIS-DMN | 180223 | -7.92 | 0.22 | **<0.0001** | - |
| SM-DA | 115236 | 0.91 | 0.03 | 10.1485 | + |
| SM-VA | 62910 | -2.11 | 0.08 | 0.9726 | - |
| SM-L | 10202 | 1.77 | 0.08 | 2.1694 | + |
| SM-FP | 128883 | 0.36 | 0.02 | 20.1562 | + |
| SM-DMN | 216799 | -8.03 | 0.2 | **<0.0001** | - |
| DA-VA | 47254 | -1.42 | 0.07 | 4.3885 | - |
| DA-L | 4284 | 0.22 | 0 | 23.2316 | + |
| DA-FP | 96899 | 2.66 | 0.09 | 0.2193 | + |
| DA-DMN | 171373 | -4.07 | 0.11 | **0.0013** | - |
| VA-L | 2543 | -2.08 | 0.13 | 1.0592 | - |
| VA-FP | 44170 | -5.28 | 0.21 | **<0.0001** | - |
| VA-DMN | 120323 | -3.51 | 0.12 | **0.0126** | - |
| L-FP | 8744 | 1.13 | 0.04 | 7.2277 | + |
| L-DMN | 16915 | -0.77 | 0.04 | 12.3365 | - |
| FP-DMN | 328834 | 1.06 | 0.03 | 8.0796 | + |

**Table S5.3** Wilcoxon sign rank differences for all network pairs: 210s.

| Edge pair | *W* | *Z* | Cohen's *d* | *p* | Direction |
|---|---|---|---|---|---|
| VIS-VIS | 35391 | -1.23 | 0.07 | 6.1635 | - |
| SM-SM | 68906 | 1.94 | 0.07 | 1.4558 | + |
| DA-DA | 24182 | 0.45 | 0.01 | 18.2831 | + |
| VA-VA | 7738 | -2.72 | 0.2 | 0.1832 | - |
| L-L | 576 | 0.39 | 0.05 | 19.5475 | + |
| FP-FP | 42779 | 0.98 | 0.03 | 9.1998 | + |
| DMN-DMN | 171684 | -5.89 | 0.18 | **<0.0001** | - |
| VIS-SM | 127200 | -11.59 | 0.38 | **<0.0001** | - |
| VIS-DA | 83043 | -1.81 | 0.07 | 1.9874 | - |
| VIS-VA | 30372 | -11.03 | 0.48 | **<0.0001** | - |
| VIS-L | 10582 | -1.34 | 0.06 | 5.0106 | - |
| VIS-FP | 70549 | -10.57 | 0.38 | **<0.0001** | - |
| VIS-DMN | 201549 | -8.32 | 0.24 | **<0.0001** | - |
| SM-DA | 108204 | -4.31 | 0.14 | **0.0005** | - |
| SM-VA | 71021 | -2.96 | 0.1 | 0.0874 | - |
| SM-L | 14141 | 1.84 | 0.08 | 1.8345 | + |
| SM-FP | 124544 | -1.98 | 0.06 | 1.344 | - |
| SM-DMN | 250760 | -5.07 | 0.13 | **<0.0001** | - |
| DA-VA | 56372 | -0.83 | 0.04 | 11.4223 | - |
| DA-L | 6682 | 0.27 | 0.02 | 21.9848 | + |
| DA-FP | 103792 | -2.34 | 0.08 | 0.5437 | - |
| DA-DMN | 246726 | 0.33 | 0.01 | 20.8045 | + |
| VA-L | 6117 | -0.68 | 0.03 | 13.8836 | - |
| VA-FP | 59531 | -3.98 | 0.15 | **0.0019** | - |
| VA-DMN | 155732 | -3.61 | 0.12 | **0.0085** | - |
| L-FP | 10435 | 0.47 | 0.03 | 17.8786 | + |
| L-DMN | 25526 | -1.12 | 0.04 | 7.3863 | - |
| FP-DMN | 363748 | -0.51 | 0.01 | 17.1242 | - |

**Table S5.4** Wilcoxon sign rank differences for all network pairs: 280s.

| Edge pair | *W* | *Z* | Cohen's *d* | *p* | Direction |
|---|---|---|---|---|---|
| VIS-VIS | 48931 | 3.52 | 0.13 | **0.0122** | + |
| SM-SM | 74149 | 1.31 | 0.06 | 5.3162 | + |
| DA-DA | 25818 | 1.51 | 0.08 | 3.6993 | + |
| VA-VA | 6549 | -5.3 | 0.37 | **<0.0001** | - |
| L-L | 656 | -0.07 | 0.03 | 26.5352 | - |
| FP-FP | 43364 | 0.08 | 0.01 | 26.1381 | + |
| DMN-DMN | 209570 | -3.12 | 0.08 | 0.0507 | - |
| VIS-SM | 155458 | -9 | 0.28 | **<0.0001** | - |
| VIS-DA | 114459 | 3.07 | 0.11 | 0.06 | + |
| VIS-VA | 28647 | -12.83 | 0.58 | **<0.0001** | - |
| VIS-L | 13256 | -2.03 | 0.11 | 1.1934 | - |
| VIS-FP | 105044 | -5.19 | 0.18 | **<0.0001** | - |
| VIS-DMN | 309889 | -2.01 | 0.06 | 1.2389 | - |
| SM-DA | 107435 | -5.04 | 0.17 | **<0.0001** | - |
| SM-VA | 76684 | -3.97 | 0.15 | **0.002** | - |
| SM-L | 18936 | 3.62 | 0.16 | **0.0084** | + |
| SM-FP | 114367 | -4.67 | 0.14 | **0.0001** | - |
| SM-DMN | 279969 | -5.8 | 0.15 | **<0.0001** | - |
| DA-VA | 57023 | -1.59 | 0.06 | 3.1434 | - |
| DA-L | 8984 | 1.06 | 0.08 | 8.091 | + |
| DA-FP | 101163 | -4.69 | 0.16 | **0.0001** | - |
| DA-DMN | 271213 | -0.89 | 0.03 | 10.3896 | - |
| VA-L | 5135 | -2.22 | 0.13 | 0.7335 | - |
| VA-FP | 55212 | -5.92 | 0.22 | **<0.0001** | - |
| VA-DMN | 153762 | -4.46 | 0.13 | **0.0002** | - |
| L-FP | 15840 | 2.8 | 0.15 | 0.1453 | + |
| L-DMN | 33413 | -0.08 | 0.01 | 26.1546 | - |
| FP-DMN | 374128 | -1.83 | 0.05 | 1.8676 | - |

**Table S5.5** Wilcoxon sign rank differences for all network pairs: 349s.

| Edge pair | *W* | *Z* | Cohen's *d* | *p* | Direction |
|---|---|---|---|---|---|
| VIS-VIS | 37519 | -1.36 | 0.05 | 4.86 | - |
| SM-SM | 84233 | 1.8 | 0.07 | 2.0255 | + |
| DA-DA | 30021 | 3.29 | 0.2 | **0.0283** | + |
| VA-VA | 8414 | -4.05 | 0.27 | **0.0015** | - |
| L-L | 873 | 0.37 | 0.02 | 19.9301 | + |
| FP-FP | 44586 | 0.5 | 0.01 | 17.3495 | + |
| DMN-DMN | 229661 | -1.52 | 0.05 | 3.5678 | - |
| VIS-SM | 186045 | -4.11 | 0.13 | **0.0011** | - |
| VIS-DA | 122508 | 4.18 | 0.14 | **0.0008** | + |
| VIS-VA | 36803 | -11.54 | 0.49 | **<0.0001** | - |
| VIS-L | 14286 | -1.43 | 0.08 | 4.299 | - |
| VIS-FP | 118300 | -3.85 | 0.13 | **0.0033** | - |
| VIS-DMN | 301926 | -2.76 | 0.07 | 0.1602 | - |
| SM-DA | 113275 | -5.49 | 0.18 | **<0.0001** | - |
| SM-VA | 92920 | -2.06 | 0.08 | 1.1029 | - |
| SM-L | 25535 | 5.37 | 0.26 | **<0.0001** | + |
| SM-FP | 121315 | -5.03 | 0.15 | **<0.0001** | - |
| SM-DMN | 310473 | -3.7 | 0.09 | **0.0061** | - |
| DA-VA | 70110 | 0.38 | 0.01 | 19.6068 | + |
| DA-L | 10979 | 0.75 | 0.04 | 12.7418 | + |
| DA-FP | 114554 | -2.84 | 0.1 | 0.1245 | - |
| DA-DMN | 300765 | 1.8 | 0.04 | 2.0185 | + |
| VA-L | 7402 | -0.7 | 0.04 | 13.6363 | - |
| VA-FP | 76660 | -2.54 | 0.09 | 0.3123 | - |
| VA-DMN | 196252 | -0.77 | 0.02 | 12.3001 | - |
| L-FP | 17742 | 2.04 | 0.11 | 1.165 | + |
| L-DMN | 41405 | 1.01 | 0.04 | 8.71 | + |
| FP-DMN | 429882 | 0.86 | 0.02 | 10.86 | + |

Two-tail sign rank testing of condition differences for all network combinations. Prior to testing, network ICC scores were averaged and ordered according to the seven Yeo functional networks: visual (VIS), somatomotor (SM), dorsal attention (DA), ventral attention (VA), limbic (L), frontoparietal (FP), and default mode network (DMN). P values are Bonferroni-corrected across all possible network combinations (28). Direction of change is listed as follows: +) greater stability under ayahuasca, -) lesser stability under ayahuasca.

**Permutation testing.**

To assess the statistical significance of the observed differential identifiability and success rate, we employed a permutation testing approach. For added clarification, success rate (Finn, Shen et al. 2015) is defined as the percentage of subjects whose identity was correctly predicted out of the total number of subjects. First, at each iteration of the permutation testing, subjects' test-retest connectomes were randomly shuffled, then differential identifiability and success rate were computed on the randomized identifiability matrix. This procedure is repeated 1000 times to generate a non-parametric "null" distribution of differential identifiability and success rate scores. Then, the observed (true) differential identifiability and success rate scores were then compared against their corresponding null distribution to determine the p-values (Nichols and Holmes 2002). Figure S1 shows all outcomes were significantly different to null modelled equivalents.

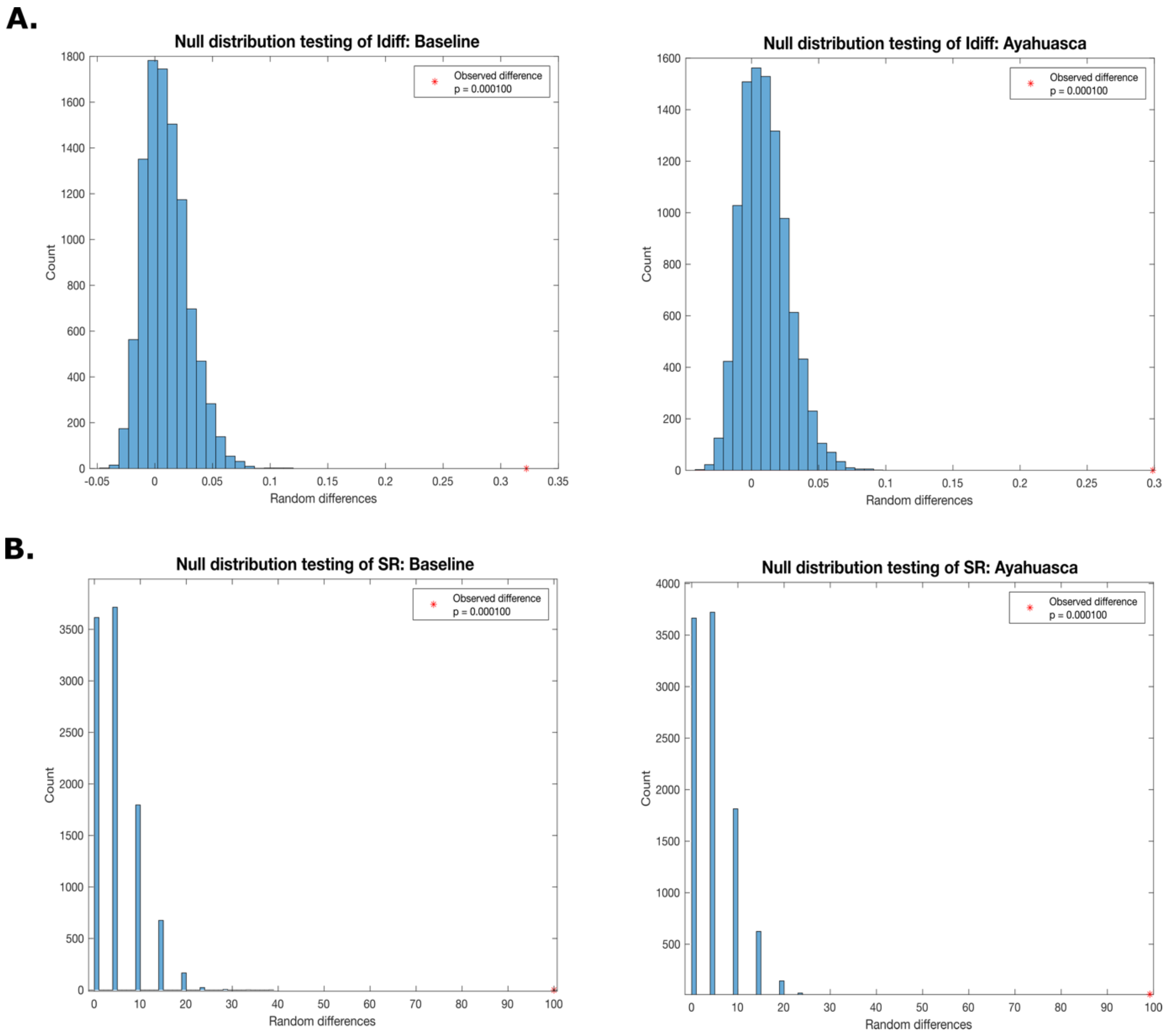


**Figure S1.** Permutation of observed $I_{diff}$ and success rate. (A) shows observed $I_{diff}$ scores (*) for each condition against their equivalent null distributions. (B) shows observed success rate $_{scores}$ (*) for each condition against their equivalent null distributions. For each panel, p-values from permutation testing are listed.

**Motion assessment.**
As a quality assessment measure we repeated our static connectome fingerprinting workflow using BOLD time series censored according to a scrubbing index devised from three different FSL-derived motion metrics: 1) Frame Displacement (FD, in mm); 2) dVARS (D referring to temporal derivative of BOLD time courses, VARS referring to root mean square variance over voxels) (Power, Mitra et al. 2014). 3) SD (standard deviation of the BOLD signal within brain voxels at every time-point). With all identifiability measures and approaches being devised from the original, static (or mean over time) FC, any significant differences arising as a result of motion would become apparent following the inclusion of this index and the removal of spurious TRs. Henceforth, the outcome *scrub/scrubbing* refers to: Total number of volumes - number of invalid (scrubbed) volumes.

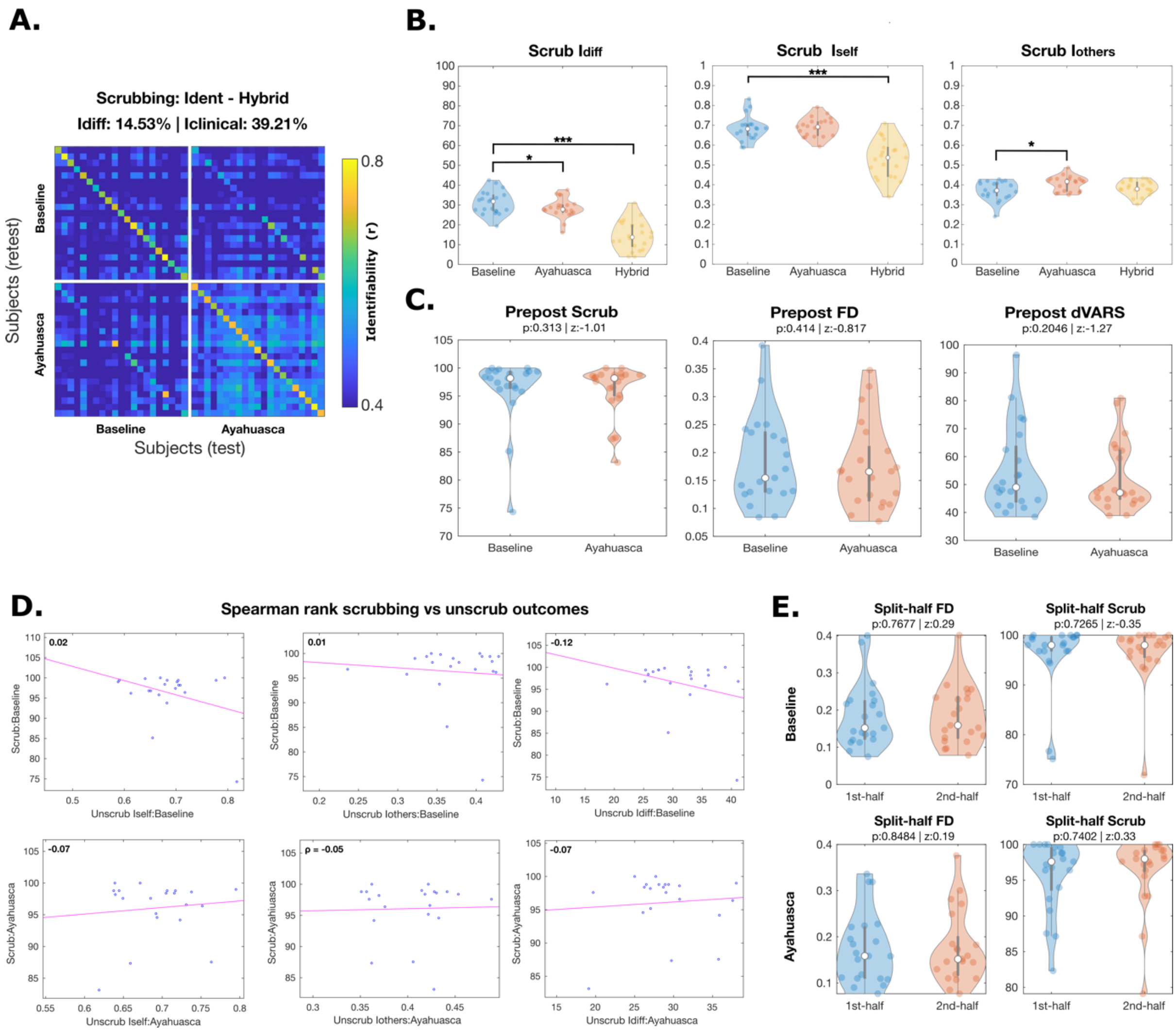

**Figure S2**. Effects of motion on identifiability differences. (A) The hybrid identifiability matrix is computed for each group, using censored test-retest individual connectomes; the resulting block matrix is composed of “standard” identification matrices (blue:baseline, orange:ayahuasca) for each condition which are expanded on the right , plus the off-block “hybrid” elements which encode the individual similarity between subjects from different conditions. In standard matrices, diagonal elements indicate self/homo-similarity ($I_{self}$) whereas off diagonal elements reflect other/hetero-similarity ($I_{others}$). From hybrid off-block elements one can also define the $I_{clinical}$ for a participant as the average similarity of the individual connectome of a subject with respect to the baseline. For all, differential identifiability ($I_{diff}$) values and success rates (SRs, where applicable) on top also provide complementary scores of the fingerprint level (see Methods). (**B**) Violin plots highlighting the difference of each identifiability metric generated using volume censoring, between conditions. Hybrid counterparts are also presented in respect to baseline. (**C**) Demonstrates the differences in scrubbing, FD, and dVARS (scrubbing is derived from the latter two) between conditions alongside *p* and approximated z-score values of sign-rank testing. **(D)** shows the association between all identifiability outcome measures and scrubbing, as assessed by Spearman rank correlation. **(E)** Highlights split-half differences in major motion outcomes, FD and scrubbing for each condition. For all violin plots, each boxplot extends from the lower to upper quartile values with a line at the median; the whiskers extend from the upper/lower quartiles up to 1.5 times the interquartile range. Subjects are represented with single points. Two-tail significance is denoted as follows: *p<0.05*, p<0.01**,p<0.001****. For all correlations, the Spearman rho is included in bold. Significant values ($p < 0.05$) are indicated by red numbering for rho.

As shown in Figure S2, no major differences were observed following the use of time series scrubbing. Note in figure S2A the increases in inter-subject correlation ($I_{others}$) under Ayahuasca, as indicated by the off-diagonal elements in the Ayahuasca matrix (bottom, right) and the diminished, intra-subject similarity across conditions ($I_{selfHybrid}$) marked by the off-diagonal matrix panels. Overall scores show minor differences to our unscrubbed findings (Scrub.$I_{diff}$ = 14.53%, $I_{clinical}$ = 39.21%; Unscrub.$I_{diff}$ 14.52%, $I_{clinical}$ = 39.21% ) however each significant outcome and direction is the same regardless of the preprocessing approach (see Fig.S2B).

We also sought to investigate whether any differences in the volumes tagged for censoring and the principal parameters determinant of this criteria might arise. Per figure S2C, no significant differences could be identified in either component across conditions. In addition, since identifiability outcomes are calculated per condition - we examined correlations between our scrubbing parameter and each of our derived, primary, unscrubbed identifiability outcomes, finding no significant associations (Fig.S2D). Since our dynamic outcomes were also derived from the same FC approach, we repeated the same approach, also finding no significant associations (Fig.S3). Note that Spearman rank correlations were employed due to the non-normality of our scrubbing vector.

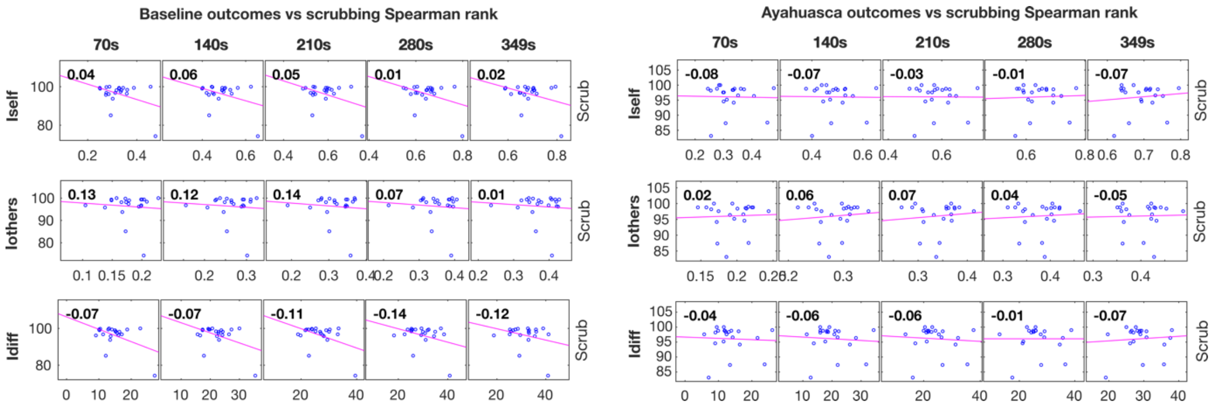


**Figure S3.** Motion correlations with dynamic identifiability outcomes. For each timescale, the association between all identifiability outcome measures and the number of volumes left after motion tagging (scrub) is assessed by Spearman rank correlation. For all correlations, the Spearman rho is included in bold. Significant values ($p < 0.05$) are indicated by red numbering for rho.

For completeness, we also evaluated motion in regard to the approach we devised to generate test-retest connectomes. In the present paper, by dividing each resting-state acquisition into halves we were able to produce test-rest scans for each condition. However, this also raises the question of how temporal differences in motion also affect identifiability. For example, it could be the case that lower test-retest correlations could arise as a result of subject's head motion varying across runs. Here, as exemplified in figure S2E, we show no within-condition differences in volumes tagged for scrubbing. In the same subpanel, we also examined framewise displacement, it being the primary covariate of interest for all prior fingerprinting work, and identified no significant differences.

**Edge standard deviation**.

As a secondary analysis, we also sought to examine whether any changes in edge stability may be mirrored by changes in a measure of signal variance such as standard deviation. A drop in edge stability may also imply greater FC variance. As per figure S4 we calculated the average test-retest standard deviation of edges for each condition. No discernible parallels could be observed, as for example SM and VIS edges exhibiting lesser variance were also seen to exhibit greater stability.

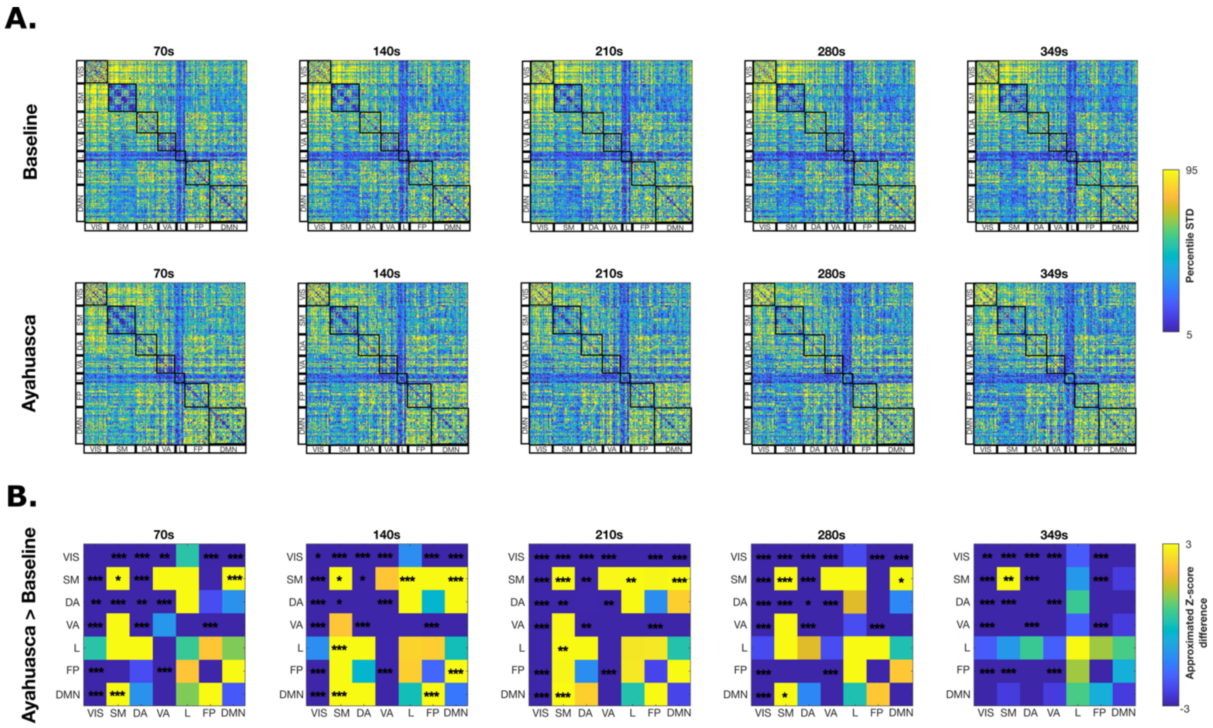


**Figure S4. Standard deviation (SD) of functional connectivity at increasing timescales.** Edgewise SD across subjects at each temporal scale for each condition. The matrices are ordered according to the seven Yeo functional networks: visual (VIS), somatomotor (SM), dorsal attention (DA), ventral attention (VA), limbic (L), frontoparietal (FP), and default mode network (DMN). (B) Differences in network SD across timescales. For each condition and per window, SD edgewise scores are averaged across Yeo functional networks and compared using bonferroni-corrected two-tail sign-rank testing. Approximated z-scores are then extrapolated and plotted for ease of visualisation.

While functional connectivity assessments of psychedelic effects have reported changes to signal complexity and variability(Carhart-Harris and Friston 2019), these measures should not be expected to be affected equally. For order statistics of non-normal distributions such as those at hand; two measures of uncertainty; standard deviation, and entropy do not convey the same information, i.e., they are not positively correlated. This is probably because the lowest and/or highest order statistics do not show high variability due to censorisation. In instances, with normal distributions, entropy and standard deviation are positively correlated as expected a priori.

**Multilinear model edge selection.**

We performed an iterative approach to derive the best performing model presented in the main manuscript. As seen from figure S5, an optimal cut-off could be defined according to when true model performance surpassed null model performance standard deviation.

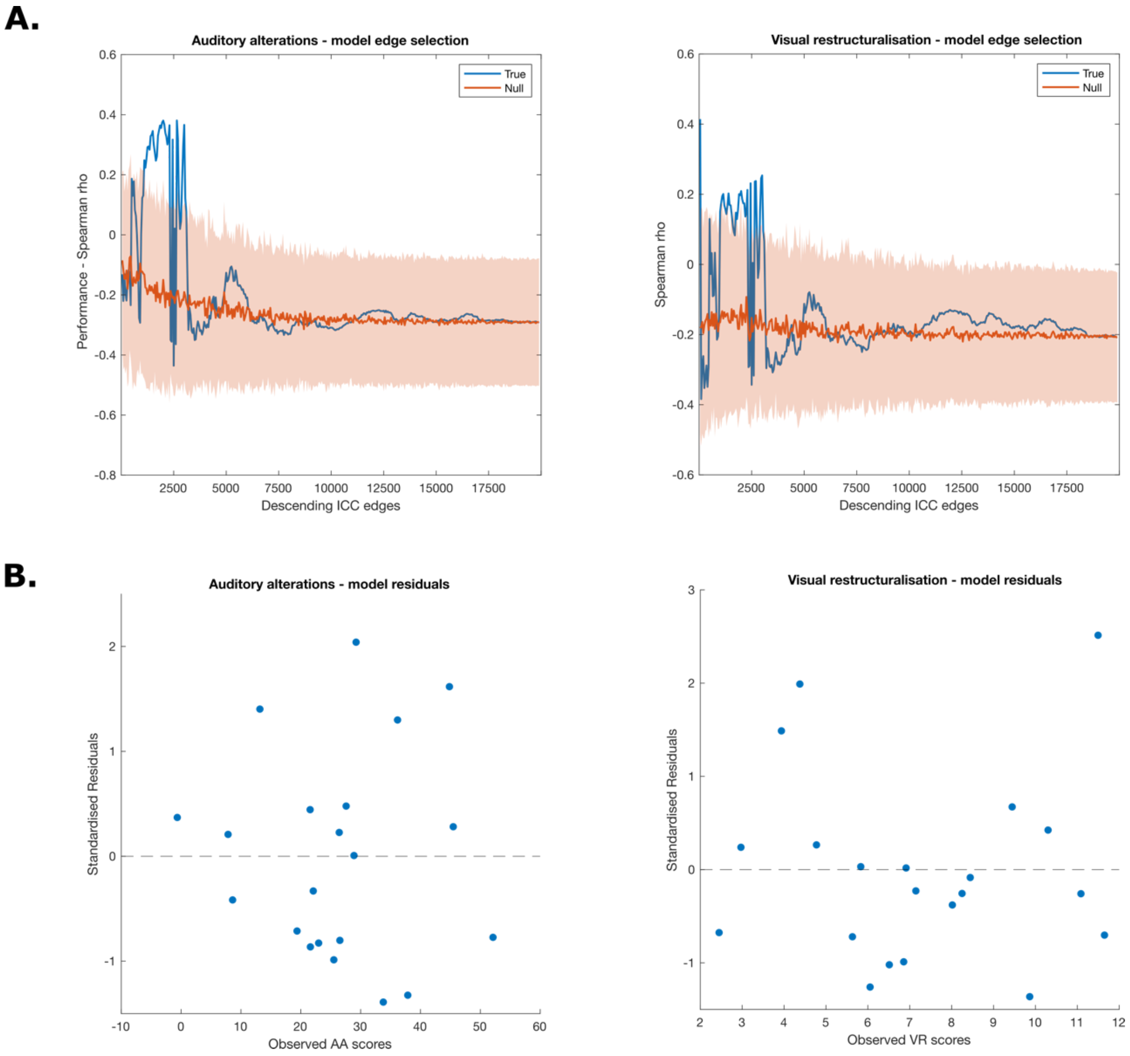


**Fig. S5. Optimal model selection. A)** Feature selection based on ICC. For ayahuasca, subset of edges are added iteratively (from 50, 100 to whole-brain, in steps of 50) based on their ICC values, from most to least reliable (x-axis), and prediction performance (k-fold (k = 5) cross validation, see Methods) of the multi-Linear model based on the PCA decomposition of selected edges and nuisance variables (PC1/2/3,singing,scrubbing) is evaluated (y-axis), and compared against a null models one (Null, red line), obtained by randomly choosing the subset edges at each step (shaded red ).
